# Metagenomic-scale analysis of the predicted protein structure universe

**DOI:** 10.1101/2025.04.23.650224

**Authors:** Jingi Yeo, Yewon Han, Nicola Bordin, Andy M. Lau, Shaun M. Kandathil, Hyunbin Kim, Eli Levy Karin, Milot Mirdita, David T. Jones, Christine Orengo, Martin Steinegger

## Abstract

Protein structure prediction breakthroughs, AlphaFold2 and ESMfold, have led to an unprecedented influx of computationally derived structures. The AlphaFold Protein Structure Database provides over 200 million predictions, while the ESM Metagenomic Atlas (ESMatlas) includes over 600 million predictions from uncultured microbes. We combine these into AFESM, an 820-million-entry dataset, and cluster them using a scalable pipeline based on sequence and structure similarity, yielding 5.12 million non-singleton structural clusters. We identify common ancestors and biomes for these clusters to explore their environmental diversity and specificity, and we investigate their structural novelties. From non-singleton clusters unique to ESMatlas, we identified 12 novel domain folds, repredicting a subset (∼45%) of low-quality domains with ColabFold yielded 33 additional novel folds. This underscores the importance of prediction quality in structural novelty discovery. We also identified 11,941 previously unseen domain combinations, highlighting the untapped structural diversity and importance of metagenomic data for illuminating underexplored regions of the protein structural universe.

An interactive webserver and data are available at afesm.foldseek.com.

## Introduction

Advancements in protein structure prediction methods, notably AlphaFold2 (Jumper et al. 2021) and ESMFold (Lin et al. 2023), have led to an unprecedented expansion of available protein structural data. The AlphaFold Protein Structure Database (AFDB) (Varadi et al. 2024) now contains over 214 million highly accurate protein structure models predicted by AlphaFold2, while ESMatlas contributes more than 600 million ESMFold-predicted structures of proteins derived from metagenomic assemblies in the MGnify database (Richardson et al. 2023). This dramatic expansion of structural data has revealed unexpected evolutionary relationships (Barrio-Hernandez et al. 2023) and highlighted novel protein domains (Lau et al. 2024) that extend our functional and evolutionary view of the protein universe. Yet, the potential of ESMatlas remains largely underexplored.

Organising and interpreting such a volume of data requires efficient computational tools. Recent algorithmic advancements, such as Foldseek (van Kempen et al. 2024), have enabled the rapid clustering of proteins based on structural similarity (Barrio-Hernandez et al. 2023). Using Foldseek, AFDB could be grouped into approximately 2.3 million distinct structural clusters. This clustering not only reduced redundancy but also uncovered intriguing structural relationships, including previously unknown connections between components of the human innate immune system and bacterial proteins.

Another large-scale effort has been made to segment AFDB into constituent domains, which are the fundamental units of protein structure and function, and organise them in the Encyclopedia of Domains (TED) (Lau et al. 2024). TED identified nearly 365 million domains across more than a million taxa. This work also included over 100 million domains previously undetected by sequence-based methods and reveals that 77% of non-redundant domains are similar to existing domains classified in the CATH database (Sillitoe et al. 2021), demonstrating the effectiveness of structure-based approaches. Existing superfamilies, particularly those with few experimentally derived structures, are greatly expanded by AFDB domain models. Additionally, domains with no similarity to structures deposited in the Protein Data Bank (PDB) and other major domain structure classification resources unveiled over 7,000 putative novel folds across the tree of life.

Large-scale clustering and domain-dissection approaches have already highlighted the breadth of structural diversity in AFDB. Notably, a recent integrative analysis revealed a shared functional landscape across AFDB clusters, the 36 million (∼6%) high-quality subset of ESMatlas, and the Microbiome Immunity Project <u>(Szczerbiak et al. 2024)</u>.

Unlike AFDB, which primarily focuses on protein sequences from culturable and well-characterised organisms, ESMatlas derives its data from uncultured metagenomic sequences, capturing the vast, untapped ‘dark matter’ of the protein universe (Vanni et al. 2022). This dataset represents a unique opportunity to explore novel protein structures and domain combinations that are absent in existing structural databases and also measure their abundance in nature. We hypothesised that ESMatlas will greatly expand our knowledge of protein structural diversity, offering insights into unexplored regions of the protein fold space and evolutionary processes.

To investigate the largely uncharted potential of these metagenomic protein structures, we compiled a joint resource of 820 million predicted models from AFDB and ESMatlas, then performed large-scale sequence and structural clustering using MMseqs2 (Steinegger and Söding 2017, 2018) and Foldseek. We annotated these clusters by identifying putative domains using CATH, assigning taxonomic labels, and integrating biome (environmental) metadata. This allowed us not only to expand the known protein structural repertoire with metagenomic proteins but also to quantify their distribution across diverse taxa and highlight structures potentially adapted to specific habitats. Moreover, it enabled us to conduct a systematic dissection of structural novelty, from uncovering genuinely new folds to pinpointing novel combinations of domains, thereby offering deeper insights into the evolutionary processes that shape protein architecture across diverse ecosystems.

## Results

### Clustering 820 million predicted protein structures

We first constructed a computational pipeline consisting of two-step clustering and various taxonomic and functional analyses (see Methods, Fig. 1) and applied it to the combined set of 820 million entries from AFDB and ESMatlas, henceforth referred to as “AFESM”.

**Figure 1.**
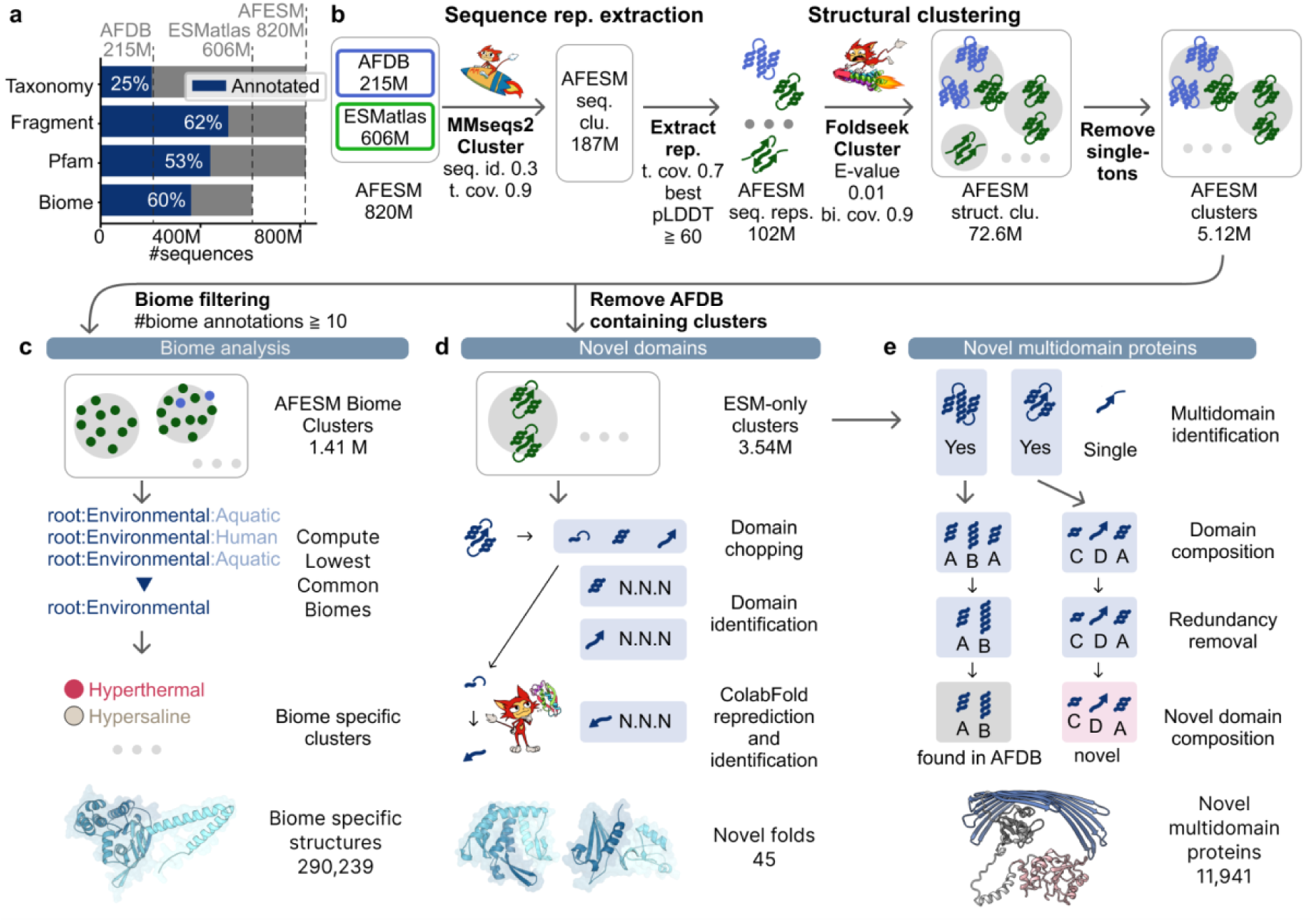
Overview of database, clustering steps and branches of analysis. a, Distribution of the metadata over AFDB, ESMatlas and their concatenation in this study: AFESM. Fractions of AFESM with various annotations are indicated in blue (Biome annotations are available for ESMatlas only) b, Clustering pipeline. AFESM was clustered using an above the twilight zone sequence identity (seq. id.) cutoff and a target to query coverage (t. cov.) that associates fragments with longer representatives. Cluster representatives were selected as the protein with the highest predicted Local Distance Difference Test (pLDDT) that is also longer than 70% of the longest protein in the cluster, discarding clusters consisting of only poor predictions (pLDDT < 60). These representatives were clustered by their structures using Foldseek Cluster, applying E-value and bi-directional coverage (bi. cov.) thresholds. c, We computed the lowest common biomes (LCB) of 1.41 million AFESM clusters with least 10 biome annotations to identify biome-specific protein structures. d, We segmented the representatives of the clusters with no AFDB entries among their members (ESM-only clusters), into putative domain regions, identified domains, and characterised their novelty. e, We identified proteins with multiple domains and characterised their domain composition. Comparing the composition against the domain arrangements in AFDB, we identified novel domain co-occurrences in ESM-only clusters.

To reduce redundancy among proteins with similar sequences and to remove protein fragments (81% of ESMatlas; fragment labels provided by MGnify (Richardson et al. 2023)), we performed sequence-based clustering using MMseqs2 Cluster (Steinegger and Söding 2018) to group fragments with their corresponding full-length proteins, if available. This clustering resulted in 187 million clusters, with the longest protein in each cluster proposed as the representative by MMseqs2. Focusing on structural information, we substituted an MMseqs2-proposed representative with a member protein if it satisfied two conditions: (1) its length was at least 70% of that of the representative, and (2) it had the highest predicted Local Distance Difference Test (pLDDT) score among all cluster members that met condition 1. Clusters represented by proteins with a pLDDT value lower than 60 were excluded, resulting in a set of 102 million high-quality representative proteins.

To discover protein similarities within the twilight zone (Rost 1999), we performed structure-based clustering on these representative proteins using Foldseek Cluster (van Kempen et al. 2024; Barrio-Hernandez et al. 2023) (see Methods). This effort resulted in 72.6 million AFESM structural clusters. Of these, 5.12 million were non-singleton AFESM clusters, much less frequent than the fraction of non-singleton clusters in AFDB (Barrio-Hernandez et al. 2023) (7% and 27% non-singleton clusters in AFESM and AFDB, respectively).

We then assessed the quality of the 5.12 million non-singleton clusters. We found that the median values of lDDT and TM-scores of member-to-representative alignments were 0.73 and 0.70, respectively (Fig. 2a), indicating generally high structural similarity within clusters. Additionally, due to the removal of low pLDDT structures and singletons during clustering, the percentage of entries with Pfam annotations increased from 53% to 70%, before and after clustering, while the percentage of fragments decreased from 62% to 55% (Fig. 1a vs. 2c).

**Figure 2.**
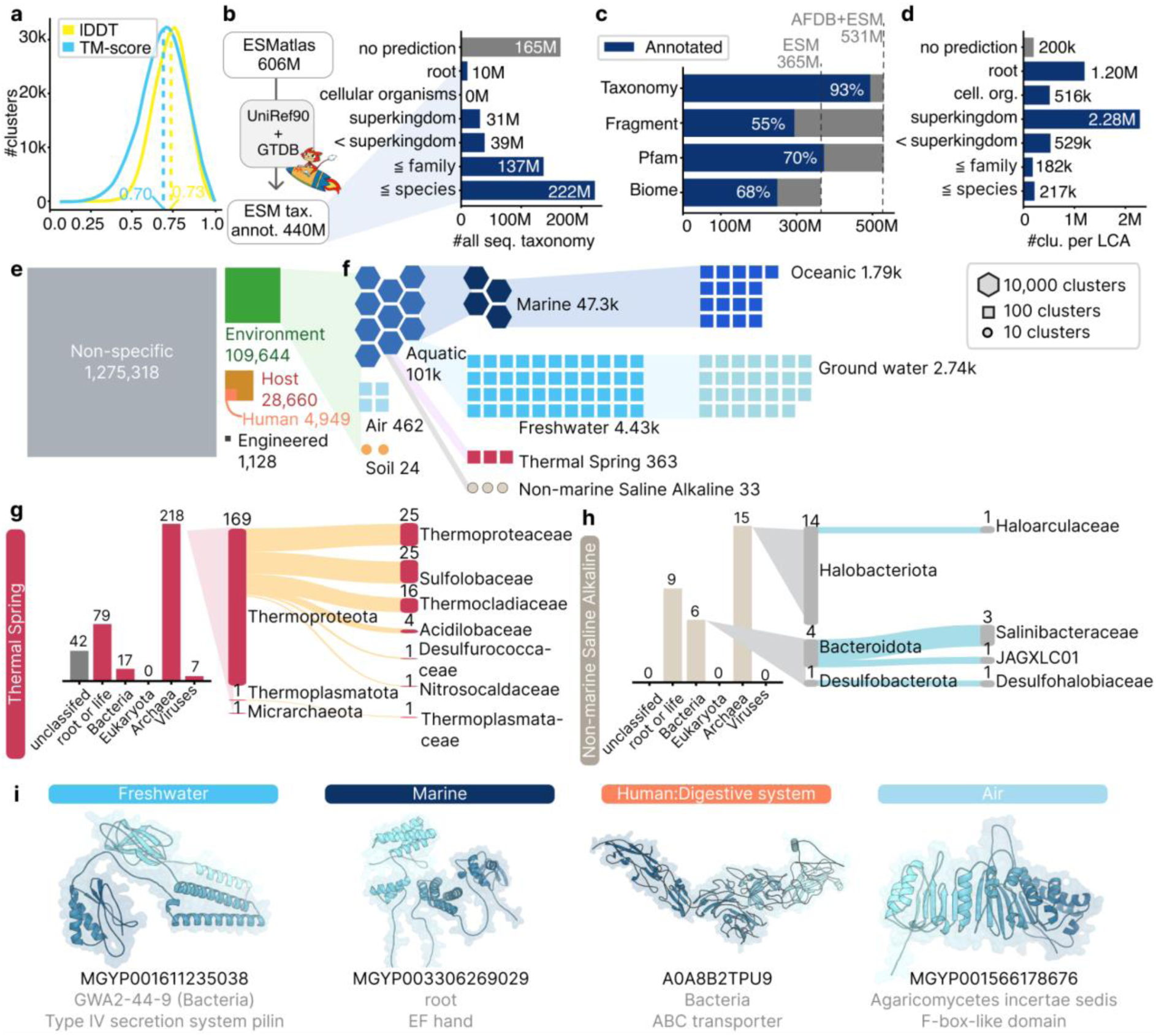
Structural similarity within AFESM clusters, their associated and projected metadata and taxonomic and biome analysis. a, Distribution of structural similarity within AFESM clusters measured as the average over all member-to-representative TM and lDDT scores. b, Millions of unannotated ESMatlas entries were assigned taxonomic labels in this study primarily to the species level. c, Distribution of the metadata over the members of AFESM clusters after projection of labels. d, LCA computation labels of AFESM clusters at various taxonomic ranks, predominantly at the superkingdom level. e, LCB labels computed for the 1.41 million AFESM biome clusters covering 238 million annotations (see Results) indicate 7.75% are environmental (green). f, Further breakdown of the environmental labels reveals clusters unique to extreme environments: Thermal Spring and Non-marine Saline Alkaline g-h, Examination of the taxonomic labels for these clusters asserts the presence of clades known to be associated with extreme environments, such as archaeal Thermoproteota for Hyperthermal-associated clusters, i, Protein examples having the most frequent Pfam in biome specific clusters.

### Structural Diversity Across Taxa

Next, we explored structural diversity across different taxa. Since ESMatlas does not include taxonomic information, we predicted taxonomic labels for the sequences by searching the ESMatlas sequences against UniRef90 (UniProt Consortium 2025) and the Genome Taxonomy Database (GTDB) (Parks et al. 2022) using MMseqs2 taxonomy (Mirdita et al. 2021), labelling 440 million (72%) of ESMatlas. Projecting these and AFDB labels onto the clustered AFESM resulted in the taxonomic labelling of 93% of the non-singleton cluster members, compared to 25% before this effort (Fig. 1a vs. 2b-c, see Methods). Among these, 222 million annotations were resolved to species or more specific levels (Fig. 2b).

We integrated this information with species-level taxonomy annotations from AFDB and calculated the lowest common ancestor (LCA) for each cluster (Fig. 2d, Supp. Fig. 1, see Methods). A total of 23.4% (1,198,802) of the non-singleton clusters were assigned the label “root”, indicating a structure found across Viruses and at least one of the three superkingdoms of cellular organisms. Similarly, approximately half of the clusters (2,283,970) were assigned to superkingdom, encompassing sequences from a single superkingdom. Around 20% (927,603) were resolved to the ranks below superkingdom, while 4% (199,879) remained unclassified.

### Biome-specific clusters capture habitat-associated structures

To characterise biome-associated protein structures in AFESM, we mapped MGnify biome annotations onto cluster members and assigned each cluster a lowest common biome (LCB) label using the MGnify biome hierarchy (see Methods). Using MGnify, we associated 362 million proteins (44% of AFESM) with biome labels (Richardson et al. 2023). To reduce bias and increase confidence in cluster-level biome assignments, we restricted the analysis to 1.41 million non-singleton clusters with ≥10 biome annotations (“AFESM biome clusters”), accounting for 238 million (66.9%) of all available biome annotations. Most AFESM biome clusters (90.1%) were assigned to the root biome, indicating they were not specific to a single biome. The remaining 290,239 clusters mapped exclusively to environmental (7.75%), host-associated (2.0%, including 0.34% human-specific), or engineered (0.07%) biomes (Fig. 2e).

We analysed the taxonomic conservation across two extreme biomes (Fig. 2f, see Methods): thermal springs (363 clusters) and hypersaline environments (33 clusters). The LCAs of these clusters were consistent with ecological expectations: thermal spring-specific clusters were predominantly linked to the archaeal Thermoproteota phylum, whereas hypersaline-specific clusters were mainly linked to archaeal Halobacteriota and bacterial Bacteroidota (Salinibacteraceae). (Fig. 2g-h). Beyond these, we examined AFESM biome clusters from five additional non-extreme environments. These also revealed dominant taxa that aligned with their environments: Air (fungal Basidiomycota), Human digestive system (Bacillota, Bacteroidota, Actinomycetota, and Uroviricota) (Hou et al. 2022; Pavia et al. 2023), Marine habitats (Pseudomonadota) (Giovannoni et al. 2005) (Supp. Figs. 2-4). Overall, the concordance between cluster taxonomy and the known ecology of these environments asserts the biological plausibility of our biome-specific assignments.

The most abundant Pfam domains in biome-specific clusters highlight protein functions consistent with habitat-specific ecological constraints or lineage-specific biology. Freshwater-specific clusters were dominated by the Type IV secretion system pilin domain (PF18895), likely reflecting the widespread occurrence of T4SS components among freshwater-associated Patescibacteria (Supp. Fig. 5a) (Quiñonero-Coronel et al. 2025). Marine-specific clusters were characterized by EF-hand calcium-binding domains (PF13202), pointing to Ca²⁺-dependent regulatory mechanisms under marine ionic conditions (He et al. 2020), (Michiels et al. 2002). In human gut-specific clusters, ABC transporter ATP-binding domains (PF00005) predominated, underscoring the importance of transport-mediated nutrient acquisition in the competitive gut environment (Koropatkin et al. 2012). Air-specific clusters most frequently contained F-box-like domains (PF12937), likely reflecting the predominance of Agaricomycetes in air-associated samples (Supp. Fig. 5b) and the expansion of F-box gene families in this lineage (Nagy et al. 2023).

Across biomes, clusters spanned hundreds to thousands of Pfam families, indicating that biome specificity is distributed across many protein families rather than confined to a small set of functions. We therefore selected representative examples and compared them to their closest matches from non-biome-specific clusters using FoldMason (Gilchrist et al. 2026), identifying structural features present in the biome-specific representatives but absent from their best matches (Supp. Fig. 6). These features provide concrete hypotheses for biome-associated divergence that can be tested in future functional and evolutionary analyses.

### Fold novelty is sparse in the ESM Metagenomic Atlas despite its scale

To investigate domain-level structural novelty across AFESM, we built upon our existing pipeline for novel fold identification (Lau et al. 2024), adding a step to exclude domains included in TED, as these are no longer novel (Supp. Fig. 7; Methods). Ideally, this pipeline could be applied to all 820 million AFESM entries; however, doing so would impose a prohibitive computational burden. We therefore focused on 3.2 million ESM-only non-singleton clusters’ representatives (hereafter, the ‘fraction of focus’; Fig. 3a; Methods), which represent structural architectures that recur in multiple proteins, increasing our confidence that these are not prediction artefacts. Within this fraction, we identified 5.1 million domains, of which 0.18 million had no CATH hit and passed the pipeline’s quality filters (Supp. Fig. 8; Methods). Of these, 12 were identified as novel folds. Another 2.3 million domains had no CATH hit but were discarded due to low quality. We therefore re-predicted them using ColabFold. This approach rescued 0.42 million domains (∼18% of the discarded dataset; Supp. Fig. 9a) that reached pLDDT scores above 70, yielding 33 additional novel folds (Fig. 3b; Supp. Fig. 8), highlighting the importance of overcoming prediction limitations for structural novelty discovery. Further evidence in support of the importance of prediction quality was obtained when we conducted the opposite experiment: we modelled all 7,427 folds previously identified as novel and included in TED using ESMfold and noticed low confidence in the ESMfold models whereas the corresponding AlphaFold models all passed the quality filters of our pipeline (Supp. Fig. 9).

**Figure 3.**
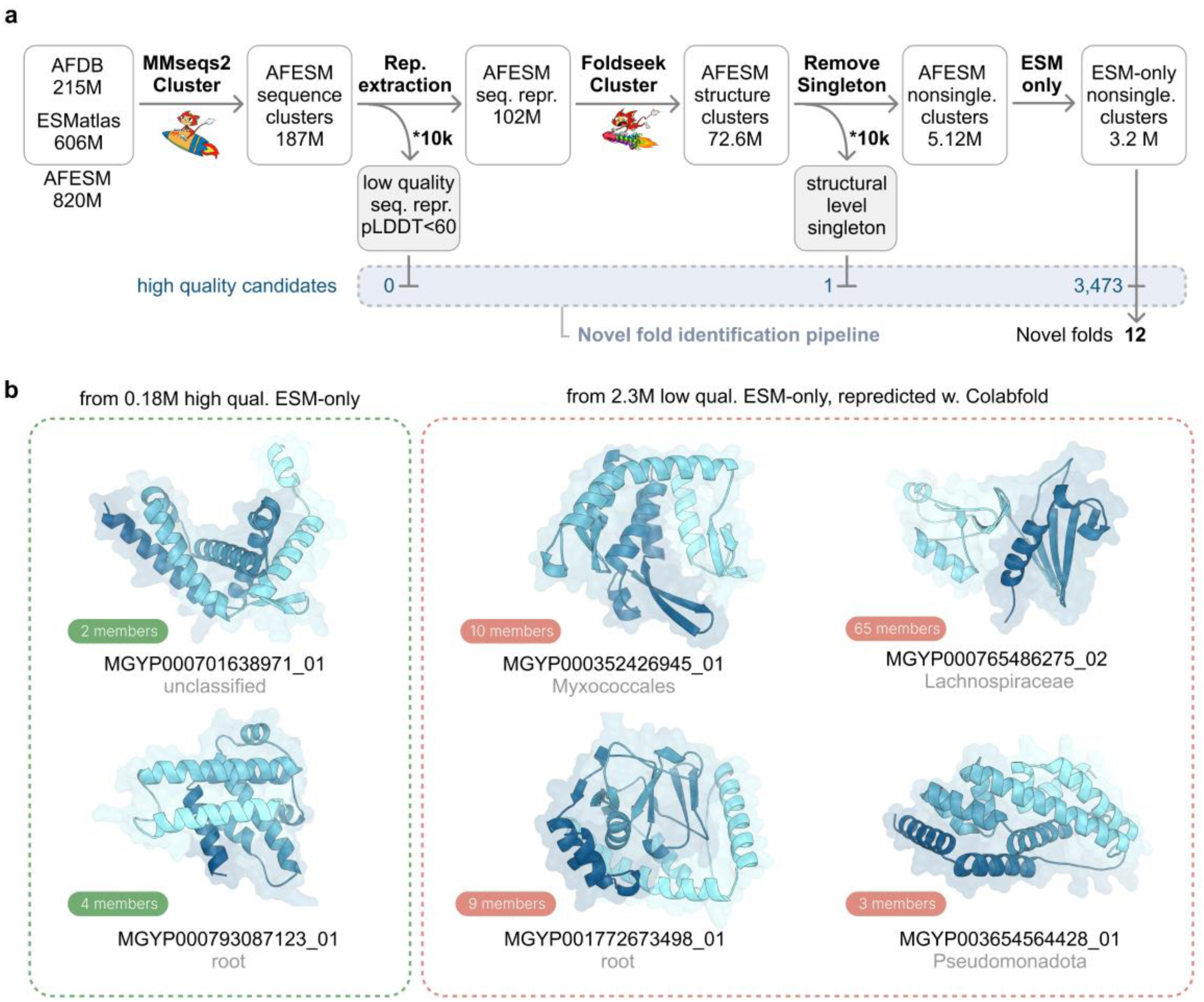
Novel fold detection within AFESM clusters. a, Schematic overview of the analysis flow used for novel fold identification. AFESM entries were sequentially filtered to define a fraction of focus consisting of ESM-only non-singleton clusters’ representatives which were analyzed by the novel domain identification pipeline. Entries excluded due to suspected low potential were additionally sampled to test this assumption (gray boxes) by counting ‘high-quality candidates’, i.e., the number of domains passing all quality filters and initial structural searches in the pipeline. The 3.2M fraction of focus yielded 3,473 high-quality candidates, from which 12 novel folds were identified based on ESMfold predictions alone. b, The two structures on the left are examples from the 12 identified novel folds. The four structures on the right are representative examples from 33 novel folds identified after ColabFold re-prediction of the 2.3 million initially low-quality, CATH-unmatched domains from the same cluster set.

Obtaining the fraction of focus was a result of applying a filtering strategy prior to the novel fold identification pipeline. Here, entries were excluded for two reasons: redundancy (sequence- or structure-level, or prior inclusion in the AFDB catalogue analysed by TED) and suspected low potential for novel fold discovery (low prediction confidence or singleton cluster status). To test whether entries discarded for low confidence (pLDDT < 60) or singleton status indeed have lower potential for fold discovery, we randomly sampled 10,000 entries from these filtering steps (Fig. 3a, gray boxes). We compared the prevalence of ‘high-quality candidates’ in the samples, i.e., domains which passed all quality filters and initial structural searches (Methods; Supp. Fig. 7; Supp. Table 1) to their prevalence in the fraction of focus. Given the pass rate of ∼0.1% in the fraction of focus (3,473 of ∼3.2 million input proteins), approximately 10 domains per 10,000 entries would be expected to reach this stage if the excluded categories had comparable novelty potential. However, we found 0 high-quality candidates in the low-confidence sample and only one in the singleton sample, supporting the assumption underlying the exclusion criteria that these categories have minimal potential.

ESMatlas is enriched relative to AFDB in short, fragmented, and non-globular proteins (Supp. Fig. 10–11), which are prone to low-confidence predictions, domain boundary ambiguity, and singleton cluster assignment—factors that likely limit detectable novelty.

However, there is reason to suspect that data quality limitations alone do not explain the sparsity of fold-level novelty in our focus fraction; 45 novel folds, including ColabFold rescues, compared to 7,427 in TED. This is further supported by independent observations: a recent large-scale exploration of nearly 1 billion metagenomic-unique proteins identified only 223 putative novel folds not present in SCOPe or PDB (Pavlopoulos et al. 2023). Furthermore, we queried the 223 structures against AFDB50 using Foldseek and found that 98% matched an AFDB entry with TM-score > 0.5, suggesting structural similarity to entries already present in AFDB. Together, these findings indicate that the discoverable fold space for recurrent proteins is approaching saturation, suggesting a common ‘domain toolbox’ shared across cellular life and metagenomes that is largely already captured in AFDB.

### Novel combinations of known domains are discovered in the ESM Metagenomic Atlas

Although 3.54 million representative structures are unique to ESMatlas (ESM-only), the few structural novelties we found cannot explain this finding. We therefore assessed how many of these unique proteins arise from previously unobserved combinations of established CATH domains rather than entirely new domains. For this we investigated the multi-domain protein (MDP) architectures of these representatives, focusing on 393,793 proteins with at least two different Topology level (T-level) CATH categories (Fig. 4a, See Methods). In order to assess how many of these harbour a CATH combination that is unique to ESMatlas, we compared them to 51.7 million proteins from AFDB that matched this criterion (Fig. 4a). This revealed 11,941 clusters with MDP architectures containing T-level CATH combinations that are unique to the ESM-only set (Fig. 4b, See Methods). When examining the novel MDP architectures, we focused on a subset of 5,203 “Novel” MDP architectures containing ≥2 distinct H-level CATH categories, the most granular CATH classification, assigned to globular domains (see Methods). Using the same criteria, we also identified in the ESM-only set 134,576 “Non-novel” H-level architectures whose domain combinations can be found in AFDB.

**Figure 4.**
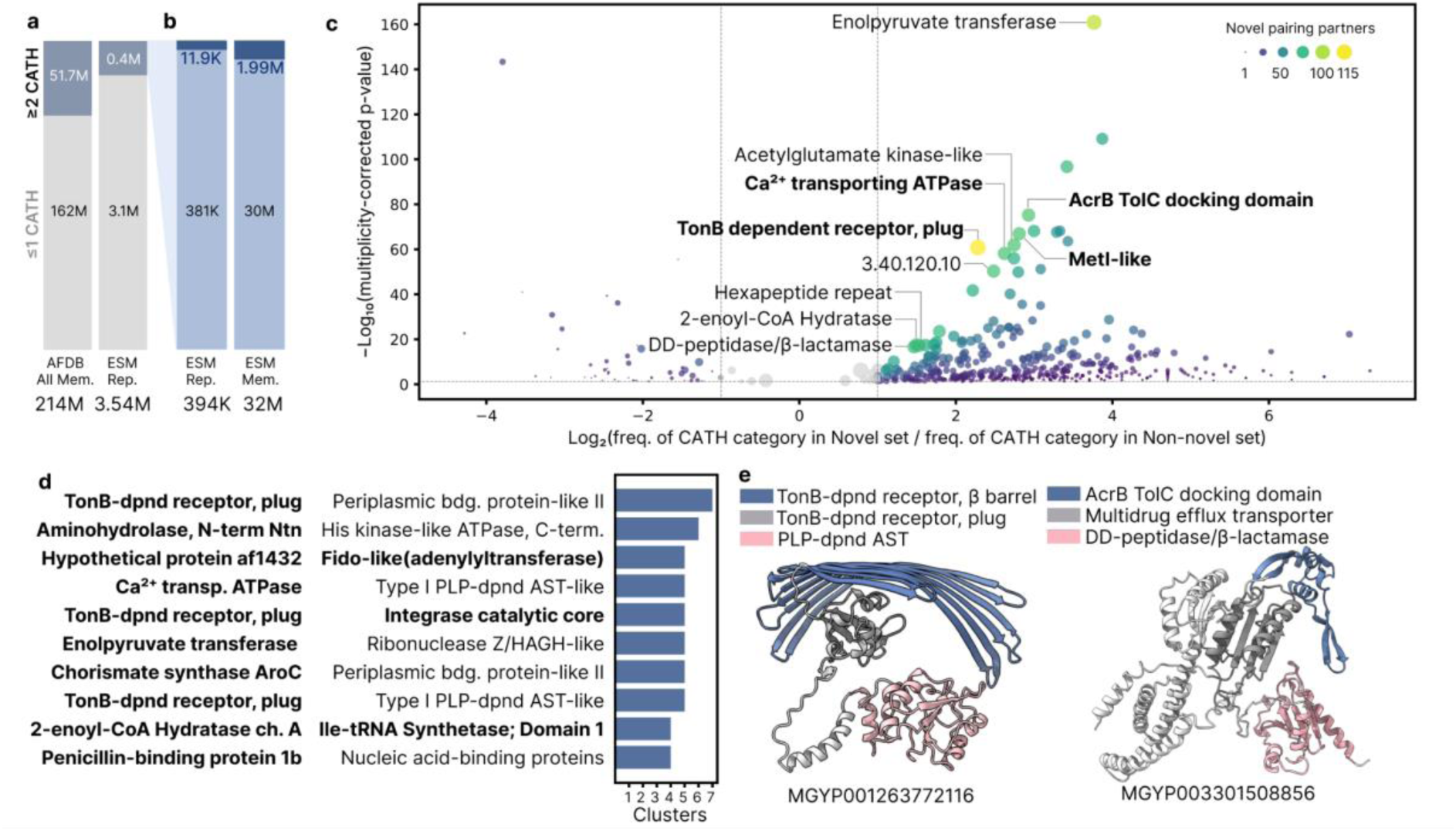
Domain combinations of ESM-only multi-domain proteins (MDPs). a, Fraction of MDPs: structures with ≥2 non-identical Topology-level (T-level) CATH annotations (blue) among AFDB members (left) and ESM-only representatives (right). b, Fraction of MDPs with novel (dark blue) co-occurring T-level CATH domains within ESM-only representatives (left) and members (right). c, Volcano plot of 530 H-level CATH domains that showed significant over/underabundance (see Methods) between ESM-only MDPs with novel domain combinations (“Novel set”) and all other ESM-only MDPs (“Non-novel set”). The x-axis indicates overabundance (right) or underabundance (left) in the Novel set and the y-axis is a function of the statistical p-value of each domain after multiplicity correction. Each domain is depicted as a circle whose colour and size reflect the number of its unique novel pairing partners, with textual labels for the top 10 and bold font for membrane-associated domains. d, Top 10 co-occurring H-level CATH domain pairs in the Novel set. Domains found to be overrepresented in the Novel set compared to the Non-novel set in panel c are highlighted in bold; the rest didn’t show a significant difference. e, Examples of novel combinations of membrane-associated CATH labels.

The joint set of 139,779 MDP architectures consisted of 2,906 unique H-level categories: 1,233 were found both in the Novel set and in the Non-novel set, 33 were unique to the Novel set, and 1,635 were unique to the Non-novel set. That is, despite being over 20× smaller (5,203 vs. 134,576 MDPs), the Novel set captured 44% (1 - (1,635/2,906)) of the categories, indicating greater domain diversity. To further characterise the Novel set, we analysed the enrichment of CATH categories between the two sets. We tested the prevalence of the 2,906 H-level categories in the two sets using Fisher’s exact or Chi-squared tests, applying Benjamini–Hochberg correction (see Methods). This identified 530 H-level categories that were significantly over- or underrepresented in the Novel set (Fig. 4c, Supp. Table 2). Of these, 431 (81.3%) were overrepresented in the Novel set, consistent with its broader domain repertoire relative to its size. Of the 10 significantly overrepresented H-level domains with the most pairing partners in the Novel set, four are membrane-associated: “Met I transporter”, “Multidrug efflux transporter AcrB”, “TonB dependent receptor”, and “Ca²⁺-transport ATPase Cytoplasmic domain” (Fig. 4c, Supp. Table 3, See Methods). Furthermore, H-level categories that are most prevalent in CATH are underrepresented in the Novel set (Supp. Fig. 12), suggesting that the discovery of combinations among well-studied domains is nearing saturation. In contrast, novel combinations of domains predominantly involve least frequent H-level categories in CATH, highlighting the potential of this analysis to uncover functional combinations involving underexplored domains.

All ten most frequently co-occurring domain pairs in the Novel set include at least one domain category that is significantly overrepresented in the Novel set (Fig. 4d). We observed pairings that appear unexpected—for instance, an outer membrane component paired with an integrase catalytic core domain, or with an aminotransferase-like domain. Unexpected domain pairings were also found in pairs involving the most overrepresented domains (Supp. Table 4), such as combinations of cytosolic and nuclear domains, or cytochrome oxidoreductase and urease peptidase. One example for a protein with unexpected domain pairing is MGYP001263772116 (Fig. 4e left). This protein contains TonB beta-barrel domains and a PLP-dependent aminotransferase domain at the N-terminus—the former typically associated with the outer membrane and the latter with the cytosol, based on PDB entries linked to their respective CATH domains. MGYP001263772116 also includes a TonB plug domain. Such a multi-domain architecture deviates from the conventional TonB-dependent transporter structure (Noinaj et al. 2010), where the N-terminal TonB box is replaced by the aminotransferase-like domain, which may be an alternative mechanism for transport regulation. Another example is MGYP003301508856, which consists of Multidrug efflux transporter beta barrel domain and DD-peptidase/β-lactamase (Fig. 4e right). This novel pairing was observed twice, suggesting a possible mechanism for hydrolysis localised at membranes.

Our MDP analysis suggests that novel combinations of known CATH domains drive the functional diversification of many metagenomic proteins.

### Limitations

The AFESM dataset contains a high fraction of fragments (56%), mostly from ESMatlas (81%). Clustering fragmented proteins by sequence and structure can miss entries which could be aligned as a member at its full length, leading to high numbers of singletons and misassigned members.

In taxonomy prediction, we observed discrepancies between reference databases. For Bacteria and Archaea, we used GTDB, a genome-based taxonomy database, whereas for Eukaryota and Viruses, we used UniRef90 (a protein database). These database differences likely contributed to the multi-modality in protein count distributions in Eukaryota and Viruses, a pattern absent in Bacteria and Archaea. Metagenomic sampling differences also introduced biases, particularly for Eukaryota and Viruses. For the former, the median of just two protein structures per genus likely underestimates true diversity due to a metagenomic sampling bias toward small organisms, whereas for viruses, these same biases underscore the difficulty of accurately assigning viral sequences to taxonomic groups.

The biome lineages defined by our LCB approach are less clearly delineated compared to taxonomic hierarchies, largely due to limitations in the biome annotation system (e.g., “Hydrothermal vents” and “Thermal springs” are not within the same lineage). We expect that adopting the recent studies on the relationship and distance between biomes based on taxonomy distribution could help to construct a reasonable lineage of the biome system (Kim et al. 2026). Additionally, many proteins in ESMatlas lack consistent or complete biome metadata and some biomes have strong sampling biases such as marines or humans, further hindering the discovery of structures specific to particular environments.

In the MDP analysis, novel protein structures comprised only 0.3% of the ESM-only clusters and 2.3% of their 820 million members (Fig. 4b). This analysis is limited by incomplete CATH coverage, excluding MDPs with fewer than two CATH annotations. In our analysis, we did not consider domain order, copy number, and spacing between domains, thus CATH superfamily names may not fully reflect domain function in multi-domain contexts.

## Discussion

In this study, we analyzed 820 million predicted protein structures, to generate AFESM, a combined and clustered resource of AFDB and ESMatlas. AFESM defines 5.12 million non-singleton structural clusters, consisting of 64.6% of the 820 million protein structures - the largest integrated predicted structure dataset to date. Most of the clusters in our analysis were assigned at the level of a superkingdom or above (78%, Fig. 1) and were not specific to a single environment (83.5%, Fig. 2), which indicates taxonomic conservation of structures across broad evolutionary lineages and habitats. Leveraging comprehensive domain annotations from the TED pipeline, we identified novel domains and further refined structural predictions through MSA-based reprediction, uncovering 45 novel folds. Although novel structural folds—those recurrently observed across different proteins—appear rare, possibly constrained by current protein language-model prediction methodologies, we discovered substantial novelty through previously unobserved domain combinations, identifying 11,941 unique multi-domain architectures. This underscores domain recombination, rather than fold discovery, as the primary mode of currently discoverable structural innovation captured within ESMatlas.

This contribution via novel domain combinations is dwarfed by the parts that escape discovery within the 3.54M ESM-only cluster representatives (Supp. Fig. 13). Definitive structural novelty was rare (∼0.3%, comprising 12 new folds and ∼11.9k combinations), 9.3% lacked identifiable domains, and 19% contained known domains (in known arrangements), however with potential additional unidentified regions. The majority (69%) consisted of fragments with known domains (16%) or were filtered out due to low prediction quality (53%, Supp. Fig. 8). This large pool of fragments (likely due to poor metagenomic assembly (Orellana et al. 2023) or poorly predicted structures may harbour additional structural novelty discoverable with full-length sequences or more robust prediction methods.

Given the many novel domain combinations we identified, future studies could leverage genomic context—such as gene neighborhoods—and multimer structure prediction methods like AlphaFold-Multimer (Evans et al. 2022), combined with large-scale comparative tools such as Foldseek-Multimer (Kim et al. 2025), to extend our analyses from single-chain proteins to multimeric complexes. We hypothesise that a significant fraction of the structural and functional diversity captured by metagenomic sequences may be encoded in protein multimers rather than monomers (Levy and Teichmann 2013).

In conclusion, this study represents the first comprehensive structural analysis integrating nearly one billion predicted protein structures from both under-characterised metagenomic sequences (ESMatlas) and the known protein universe (AFDB). We established a computational framework to systematically explore structural diversity at unprecedented scale, we uncovered structural adaptations specific to environmental niches, notably within extreme habitats. While entirely novel domain folds seem elusive—likely reflecting either genuine biological constraints or methodological limitations—the numerous previously unseen domain combinations we discovered emphasise the critical role of domain recombinations in evolutionary innovation. Our approach demonstrates how large-scale integration of isolated and metagenomic datasets can illuminate the vast, yet underexplored regions of the protein structural universe.

## Methods

### Preparation of the ESMatlas dataset

We obtained the ESMatlas dataset by downloading tar archives for both the v0 and v2023_02 releases using aria2c from the ESM repository (https://github.com/facebookresearch/esm). To optimise data handling, these combined archives were compressed using Foldcomp (Kim et al. 2023), a protein structure-specific compression format. During compression, entries with discontinuous protein chains were excluded via the --skip-discontinuous option. This process, applied to the merged content of both ESMatlas releases, yielded 605,776,560 protein structure entries used for the subsequent clustering analyses.

### Clustering the AFESM dataset

We performed the following steps: (1) Sequence clustering by MMseqs2 cluster with the following parameters—

> min-seq-id 0.3 --cov-mode 1 -c 0.9

(2) We initiated the selected representative sequence from each cluster by taking its longest sequence (mark its length as ‘x’). Then, we examined all cluster members whose length was >0.7x, substituting the representative with the highest pLDDT sequence among them. Then, we removed 84,463,443 clusters (151,488,016 sequences) whose representative had a pLDDT score lower than 60 (see Code Availability), leaving 102,104,329 representatives.

(3) Structure clustering of the 102,104,329 entries using Foldseek cluster with the following parameters

> -c 0.9

(4) During the clustering workflow, we repeatedly re-clustered representative protein structures. Because cluster representatives from one iteration become input to the next, this can result in cluster memberships that violate our defined thresholds (e.g. coverage or e-value). After the initial clustering, 38,386,513 entries formed non-singleton clusters, but 10,355,205 (27%) failed to meet our parameters. We therefore introduced a reassignment step by reassigning members that violated the thresholds to more appropriate clusters or grouping them into new clusters, reducing threshold violations of members from 10.3 million to 141,686—a 98.6% improvement among 34,624,609 non-singleton members.

> --cluster-reassign 1

### Taxonomy prediction and integration

We predicted the taxonomic labels of the ESMatlas dataset using MMseqs2 Taxonomy (Mirdita et al. 2021) with a new reference database, which was generated by merging the GTDB and UniRef90 databases as follows. Using the databases command, we downloaded GTDB (Apr 24, 2024/v220), a reference for Bacteria and Archaea and UniRef90 (Nov 8, 2023), which includes sequences from the entire tree of life. By default, UniRef90 associates each sequence (cluster member) with the taxonomic identifier (taxid) of the LCA inferred for its cluster. This tends to associate sequences with the label “root”. To use more informative labels, we instead associated Uniref90 entries with the taxid of the cluster representative instead of its LCA taxid. Next, we filtered out all bacterial and archaeal sequences from UniRef90 using the command filtertaxseqdb (excluding taxids 2 for Bacteria and 2157 for Archaea). Then, we combined GTDB with the filtered UniRef90, resulting in a merged reference containing Archaeal, Bacterial, Eukaryotic and Viral references (ABEV) and we predicted the taxids of ESMatlas by aligning against ABEV.

Meanwhile, there were remaining bacterial and archaeal sequences in AFDB with the NCBI taxonomic labels which arise because of a discrepancy of the taxonomy system between the prediction results in ESMatlas (ABEV system) and AFDB (NCBI system). We converted their labels to corresponding GTDB labels using a module provided by Metabuli (Kim and Steinegger 2024). Referring to the table offered by GTDB that describes the mapping information between GTDB taxonomic ids and NCBI taxonomic ids, we computed the taxonomic LCA of the NCBI taxonomic identifiers connected to each GTDB taxonomy due to the many-to-many pairs between the NCBI and GTDB taxonomic IDs. Using the result - one GTDB id and its NCBI taxonomic LCA, we converted the bacterial and archaeal taxonomic IDs annotated to the AlphaFold database to the corresponding GTDB identifiers.

To streamline the results, we applied the same grouping strategy, categorising the assigned taxonomy into the six hierarchical groups as root, cellular organisms, superkingdom, < superkingdom, ≦ family, ≦ species.

### Taxonomic Lowest Common Ancestor (LCA) computation

We computed the taxonomic LCA to infer the branching points of clusters using the lca module in MMseqs2. During the analysis, we excluded four taxa: (1) 12,908 unclassified sequences, (2) 28,384 sequences with “other” annotations, (3) 2 (NCBI Bacteria), and (4) 2,157 (NCBI Archaea).

### Lowest Common Biome (LCB) computation

We aimed to determine the narrowest biome region associated with each protein cluster in ESMatlas. The MGnify database defines 297 distinct biomes organised as a lineage tree that starts at the root and branches into four main categories: Environmental, Host-associated, Engineered, and Mixed. These categories further diverge into six hierarchical levels. For each cluster, we identified the LCB, representing the most specific (narrowest) biome element shared within the lineage across all cluster members. We excluded members labelled as “root” or “root:Mixed” from the analysis as their inclusion results in clusters annotated at higher-level biomes ignoring other annotations.

We computed the biome annotation on MGYP90 where ESMatlas entries were derived from. MGnify provides MGYP90 cluster information and biome annotations of the MGYP90 members. To fully utilise data, we included all members of MGYP90 clusters and annotated each ESMatlas entry with the resulting LCBs. For the AFESM Biome Clusters (see Results), we took this approach by including the biome annotations of all members within the sequence and structure clusters to calculate the LCB.

### Protein structures specific to extreme environments

We focused on three extreme environment categories: Hyperthermal, Hypersaline, and Glacier for the main analysis and five environment categories: Air, Human Digestive System, Marine for the supplementary analysis. The LCB annotations of each cluster were grouped into these categories based on their biome elements (Table 1). We examined and plotted the taxonomy distribution in each biome using Pavian (Breitwieser and Salzberg 2020).

**Table 1.** The Biome categories and their corresponding LCBs.

| Biome category | Lowest Common Biomes |
| --- | --- |
| Hyperthermal | root:Environmental:Aquatic:Thermal springs:Hot (42-90°C)<br>root:Environmental:Aquatic:Thermal springs |
| HyperSaline | root:Environmental:Aquatic:Non-marine Saline and Alkaline:Salt crystallizer pond<br>root:Environmental:Aquatic:Non-marine Saline and Alkaline:Saline:Microbial mats<br>root:Environmental:Aquatic:Non-marine Saline and Alkaline:Saline<br>root:Environmental:Aquatic:Non-marine Saline and Alkaline<br>root:Environmental:Aquatic:Non-marine Saline and Alkaline:Alkaline |
| Air | root:Environmental:Air |
| Human Digestive System | root:Host-associated:Human:Digestive system<br>root:Host-associated:Human:Digestive system:Large intestine:Fecal<br>root:Host-associated:Human:Digestive system:Large intestine<br>root:Host-associated:Human:Digestive system:Large intestine:Sigmoid colon |
| Marine | root:Environmental:Aquatic:Marine<br>root:Environmental:Aquatic:Marine:Coastal<br>root:Environmental:Aquatic:Marine:Hydrothermal vents<br>root:Environmental:Aquatic:Marine:Intertidal zone<br>root:Environmental:Aquatic:Marine:Intertidal zone:Coral reef<br>root:Environmental:Aquatic:Marine:Intertidal zone:Estuary<br>root:Environmental:Aquatic:Marine:Intertidal zone:Oil-contaminated<br>root:Environmental:Aquatic:Marine:Intertidal zone:Salt marsh<br>root:Environmental:Aquatic:Marine:Marginal Sea<br>root:Environmental:Aquatic:Marine:Oceanic<br>root:Environmental:Aquatic:Marine:Oceanic:Oil-contaminated<br>root:Environmental:Aquatic:Marine:Sediment |
| Freshwater | root:Environmental:Aquatic:Freshwater<br>root:Environmental:Aquatic:Freshwater:Drinking water:Delivery networks<br>root:Environmental:Aquatic:Freshwater:Groundwater<br>root:Environmental:Aquatic:Freshwater:Ice:Glacier<br>root:Environmental:Aquatic:Freshwater:Lake<br>root:Environmental:Aquatic:Freshwater:Lentic<br>root:Environmental:Aquatic:Freshwater:Lentic:Sediment |
|  | root:Environmental:Aquatic:Freshwater:Pond<br>root:Environmental:Aquatic:Freshwater:Sediment |

### Segmenting the ESMatlas chains into constituent protein domains

The ESMatlas chains underwent a consensus-based domain parsing workflow that incorporated three segmentation methods: Merizo (Lau et al. 2023), Chainsaw (Wells et al. 2024), and UniDoc (Zhu et al. 2023). Each method analysed the structure of the model and predicted domain boundaries. For UniDoc, which does not identify non-domain regions (NDRs), an initial step using Merizo was employed to filter out NDRs before domain segmentation was performed. This preprocessing step is essential, as the presence of NDRs can cause UniDoc to misinterpret entire chains as single domains. By first removing NDRs, the workflow ensured more accurate domain predictions using UniDoc.

### ESMatlas domain assignments to CATH superfamilies

Domains obtained via the consensus-based segmentation process were assigned to 31,574 existing CATH Structural Similarity Groups at 5Å (SSG5), where each SSG5 is a cluster representative for CATH domains that superpose within 5Å. The assignment was performed using Foldseek easy-search (version a435618b95ba95cbfe74dcbc8b4bd2720547d285) with the following parameters:

> --cov-mode 5 --alignment-type 2 -e 0.108662 -s 10 -c 0.366757 -a --format-output “query,target,fident,evalue,qlen,tlen,qtmscore,ttmscore,qcov,tcov“

Raw outputs were subsequently processed to retrieve assignments at the superfamily level (H-level) or fold-level (T-level) by using the following cutoffs identified in Lau et al. (Lau et al. 2024) for E-value (0.019000 and 0.108662), coverage (0.366757 and 0.786333), and TM-score (0.560000 and 0.416331), respectively.

### Domain-centred quality metrics

Domains were assessed using CATH-AlphaFlow (Waman et al. 2024), a Nextflow pipeline that calculates metrics on PDB files. These include the average pLDDT of the domain, the number of secondary structure elements (SSEs) identified by STRIDE (Heinig and Frishman 2004), and globularity metrics based on packing density and normed radius of gyration (Waman et al. 2024). Domains were deemed of sufficient quality for subsequent analyses with pLDDT > 70, pLDDT_80_>90 (at least 80% of residues have a pLDDT of 90 and above), at least six SSEs, packing density greater or equal to 10.333 and a normed radius of gyration less than 0.356.

### Iterative clustering and search

Domains that passed the quality assessment were clustered with foldseek cluster using the following parameters

> -s 4 -e 0.001 -c 0.7 --cov-mode 5 --tmscore-threshold 0.56 --min-seq-id 0.2 --seq-id-mode 1 --cluster-mode 1

And subsequently with foldseek easy-cluster:

> -c 0.8 --tmscore-threshold 0.8 --cluster-mode 1

Representatives obtained via the two clustering steps identified above were searched against PDB with

> easy-search models2 PDB result.m8 tmp -c 0.6 --cov-mode 2 --tmscore-threshold 0.5 --alignment-type 1 --format-output

> query,target,qlen,tlen,qtmscore,cigar

Results were filtered retaining domains with qtmscore greater than 0.56 and query coverage over 0.6 calculated from the cigar alignment.

Domains not meeting these criteria were searched against the Alldoms database with

> foldseek easy-search models3 alldoms result.m8 tmp -c 0.6 --tmscore-threshold 0.3 --alignment-type 1 --exhaustive-search 1 --format-output query,target,qlen,tlen,qtmscore,cigar

Valid domain matches to Alldoms required a qtmscore greater than 0.56, a cigar-computed coverage for both query and target above 0.6.

### Quality assessment with Foldclass DomQual

Domain quality was further assessed with a version of the Foldclass (Kandathil et al. 2025) neural network with an additional EGNN layer to score domains with a binary output layer, as outlined in Lau et al. Domains with a score below 0.5 were not processed further.

### Final exhaustive search and symmetry calculations

Domains with a Foldclass DomQual score over 0.5 were clustered with members of the Alldoms database using foldseek easy-cluster with the following parameters

> --tmscore-threshold 0.5 -c 0.6 --alignment-type 1

Clusters containing ESM-derived domains and Alldoms members were discarded. The remaining domains were searched exhaustively against the Alldoms database (i.e. disabling the pre-filter step in Foldseek, guaranteeing that every domain is searched against all target database members) with

> foldseek easy-search -c 0.6 --cov-mode 2 --tmscore-threshold 0.3 --alignment-type 1 --exhaustive-search 1 --format-output query,target,qlen,tlen,qtmscore,ttmscore,rmsd,cigar

The output of the search was filtered using the following criteria

> (max(qtmscore,ttmscore) > 0.5 OR rmsd < 3.0) and cigar-computed qcov > 0.6

Domains not meeting the criteria underwent one final search using Foldclass embeddings validated by TMalign. Hits meeting the following criteria were not considered novel: max(qtmscore,ttmscore) > 0.5 and cigar-computed qcov > 0.6

In order to assess internal symmetry SymD (Tai et al. 2014) was used with default parameters, labelling domains as symmetrical if the domain Z-score was greater than 9.

### Search against The Encyclopedia of Domains

In order to assess exclusivity to the ESMatlas database, we searched domains that underwent all previous steps against a Foldseek database for The Encyclopedia of Domains using the following parameters:

> -c 0.6 --tmscore-threshold 0.3 --alignment-type 1 --exhaustive-search 1 --format-output query,target,qlen,tlen,qtmscore,cigar

### Foldseek Search of Literature-Reported Novel Folds Against AFDB50

We downloaded the structure files of 223 proteins reported as novel-folds from the “supplementary data 3” of the reference (Pavlopoulos et al. 2023), and used Foldseek (easy-search) to query them against the 50% sequence identity clustered version of AFDB (AFDB50) version 4 downloaded from the AFDB cluster webserver https://afdb-cluster.steineggerlab.workers.dev/, using default parameters.

This resulted in AFDB hits with a query-TM score > 0.5 for 191 of the proteins. We then performed a more exhaustive search on the remaining 32 proteins, adding to the above command the flags --exhaustive-search and --cluster-search 1, enforcing a comparison to all cluster members and not only the representatives. Using this command, 27 additional proteins could be matched with query-TM score >= 0.5 against AFDB.

### Identifying CATH-MDPs in the ESM-only clusters and AFDB

From the set of cluster representatives of the 3.54 million ESM-only non-singleton clusters, we extracted 393,793 MDPs containing at least two CATH annotations—at either the T-level or H-level—with distinct T-level classifications.

In preparation for identifying novel domain co-occurrences in ESM-only proteins, we used the same criteria to identify MDPs in AFDB. To ensure comprehensive coverage, we used all 214 million AFDB entries (“members”) rather than just the cluster representatives. From this set, we identified 92.9 million MDPs, of which 51.7 million contained at least two CATH-annotated domains, based on annotations from the TED database (Lau et al. 2024). Additionally, we included 625K TED domains without CATH annotations but meeting high-quality standards (see ‘Domain-centered quality metrics’). These domains were searched against the CATH database using Foldseek exhaustive search with the below-specified parameters. For 25% of the 625K domains containing discontinuous regions, the starting and ending points were treated as domain boundaries.

> --exhaustive-search 1 --format-output query,target,qlen,tlen,qtmscore,cigar

For the domains identified with CATH hits through the Foldseek search results, the hits with the highest target TM scores were selected as CATH domain annotations, augmenting the TED dataset to create a comprehensive set of 51,741,054 CATH-annotated MDPs within AFDB.

### Identifying MDPs with novel CATH domain pairings

To identify domain co-occurrences unique to ESM-only proteins, 393.8K MDPs from the ESM-only, non-singleton cluster representatives were compared against 51.7M MDPs from AFDB. Each MDP was mapped to a non-redundant, order-invariant set of CATH T-level categories (e.g., an MDP with three domains: ‘1.10.10.10’, ‘1.10.10’, and ‘2.20.20.20’ maps to {1.10.10, 2.20.20}). All possible unordered domain pairs were extracted per MDP and compared to those from AFDB to identify novel pairs exclusive to the ESM-only set.

An MDP was considered “novel” if it contained at least one pair not observed in AFDB; 11.9K MDPs met this criterion (the “Novel set”), while the remaining 381.5K formed the “Non-novel set.” Novelty was assessed at the T-level rather than H-level for two reasons: (1) it maximises explanatory coverage using available data, and (2) it applies a stricter criterion, reducing potential false positives caused by inconsistencies of structural diversity within each H-level categories.

We further selected MDPs with ≥2 globular CATH H-level annotations as they are likely functional. From the 11.9K Novel MDPs, 5,203 met this criterion; from the 381.5K Non-novel set, 134,576 were selected. For example, MGYP2 ({Globular:2.20.20.20, Globular:3.30.30.30}) is included, while MGYP1 ({Globular:1.10.10.10, Non-globular:1.10.10.20, Non-globular:3.30.30}) is excluded due to having only one globular H-level domain.

### Composition analysis of the Novel MDP architectures

We compared H-level CATH category representation between the Novel (5,203 MDPs) and Non-novel (134,576 MDPs) sets. Together, the sets consisted of 2,906 unique H-level domains and the frequency of each domain in each set was computed as the number of occurrences of that domain in the set divided by the total number of MDPs in the set. Using these frequencies, we computed the differential prevalence of each domain as log₂(frequency_Novel / frequency_Non-novel).

We examined associations between the prevalence of each of the 2,906 domains and the type of set (“Novel”/”Non-novel”) as follows. In its turn, we treated each domain as ‘x’, defining all other domains as ‘not-x’. We then counted the occurrences of ‘x’ and ‘not-x’ in the Novel and Non-novel sets, constructing a 2×2 contingency table to test if the null hypothesis stating “no difference in the prevalence of x between Novel and Non-novel” can be rejected. Using the contingency table we performed a Fisher’s exact test for 2,511 of the domains and a Chi-squared test with Yates’ correction for 395 of the domains. Fisher’s test was used when low counts caused errors in the Chi-squared test.. The tests were conducted using R version 4.4.3 and the complete list of 2,906 P-values was corrected for multiple hypothesis testing using the Benjamini–Hochberg procedure (Benjamini and Hochberg 1995) with FDR < 0.05, as the cutoff.

For the top 10 by the number of their unique novel pairing partners, we examined the FunFam (Scheibenreif et al. 2019) descriptions corresponding to their CATH identifiers on the CATH website (accessed April 2025), to ensure the consistency between the CATH name and the FunFam. H-level category names were assigned using cath-names.txt (CATH v4.3.0). To determine membrane association of CATH categories, we queried the CATH web service for GO ‘Cellular Component’ terms linked to PDB entries. Categories were labelled as membrane-associated if the majority of annotations indicated membrane localisation (e.g., 87.4% of GO terms for 2.170.130.10 correspond to “outer membrane”).

## Software used for analysis

MMseqs2: MMseqs2 v14 for clustering, 407b31 for MSA generation of the domains to predict. Foldseek: ef768f for clustering. ColabFold: 1.5.5 (fdf3b235) for the domain prediction. Python 3.11: matplotlib 3.6.2 and pandas 2.2.2 for visualization, R 4.4.3, Pavian 1.2.1

## Webserver

We adapted our previously-developed AFDB clusters webserver (Barrio-Hernandez et al. 2023) to allow for exploration and visualisation of AFESM. This adaptation includes additional metadata. We will implement biome search functionality in the future.

## Data and code availability

All scripts to rerun the analysis are available at https://github.com/steineggerlab/afesm-analysis, the data can be explored and downloaded at https://afesm.foldseek.com.

## Acknowledgements

We thank Dr. Pedro Beltrao initial discussion regarding the analysis, and Dr. Robert D. Finn for insightful discussions regarding the MGnify database and for maintaining this essential resource, and Dr. Hyunbin Lee for valuable insights on the novel membrane protein examples. M.S. acknowledges support by the National Research Foundation of Korea grants (2020M3A9G7103933, RS-2021-NR061659 and RS-2021-NR056571, RS-2024-00396026), Novo Nordisk Foundation (NNF24SA0092560), Samsung DS Research Fund and Creative-Pioneering Researchers Program through Seoul National University. M.M. acknowledges support from the National Research Foundation of Korea (grant RS-2023-00250470). This work was supported by the Biotechnology and Biological Sciences Research Council (grant BB/T019409/1 to A.M.L. and D.T.J., grant BB/W008556/1 to S.M.K. and D.T.J.) and the Wellcome Trust (grant 221327/Z/20/Z for N.B. and C.O).

## Declaration of interests

M.S. declares an outside interest in Stylus Medicine as a scientific advisor.

## Supplementary materials and methods

**Supplementary Figure 1.**
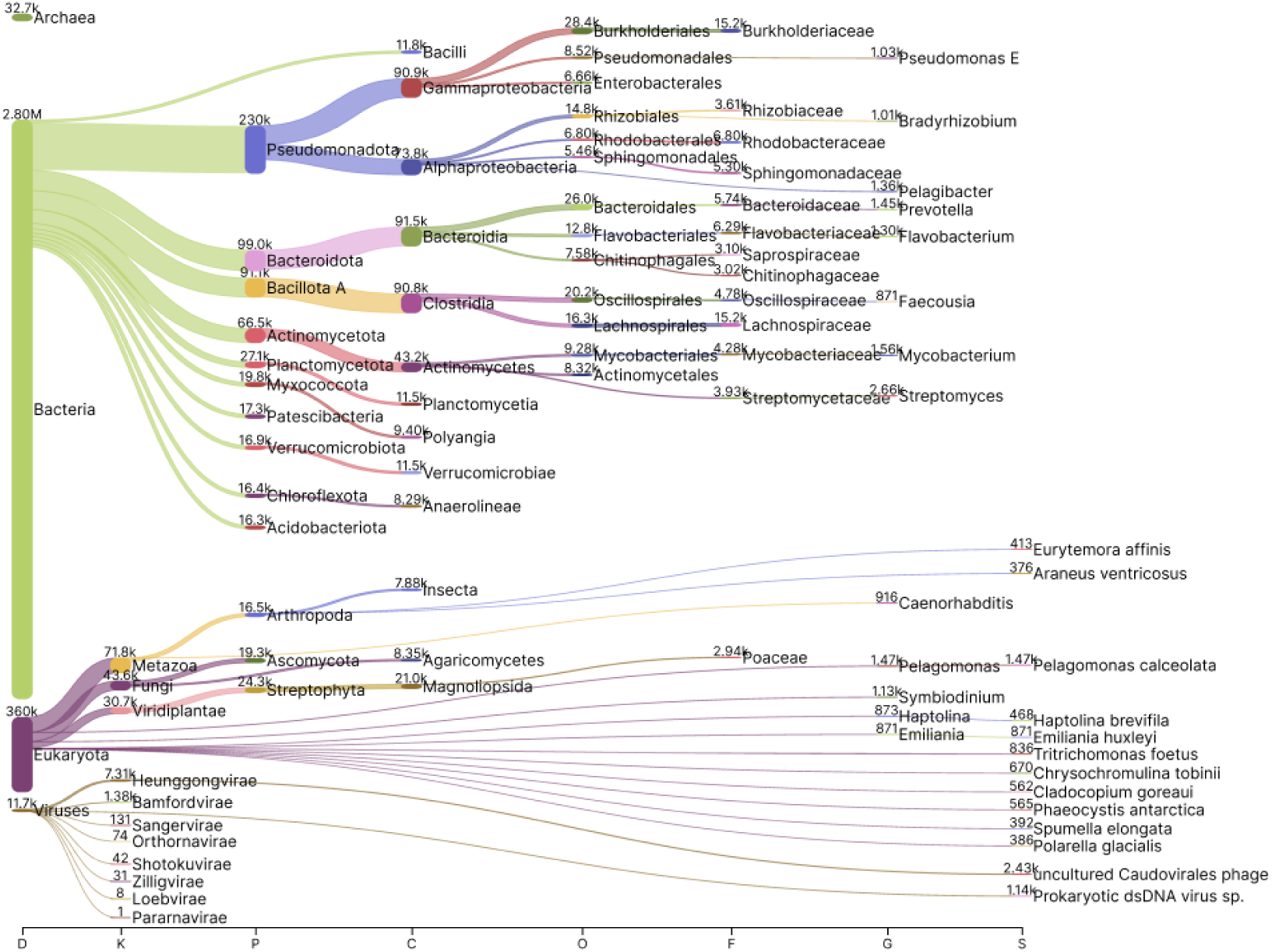
The taxonomic distribution of the non-singleton AFESM clusters.

**Supplementary Figure 2.**
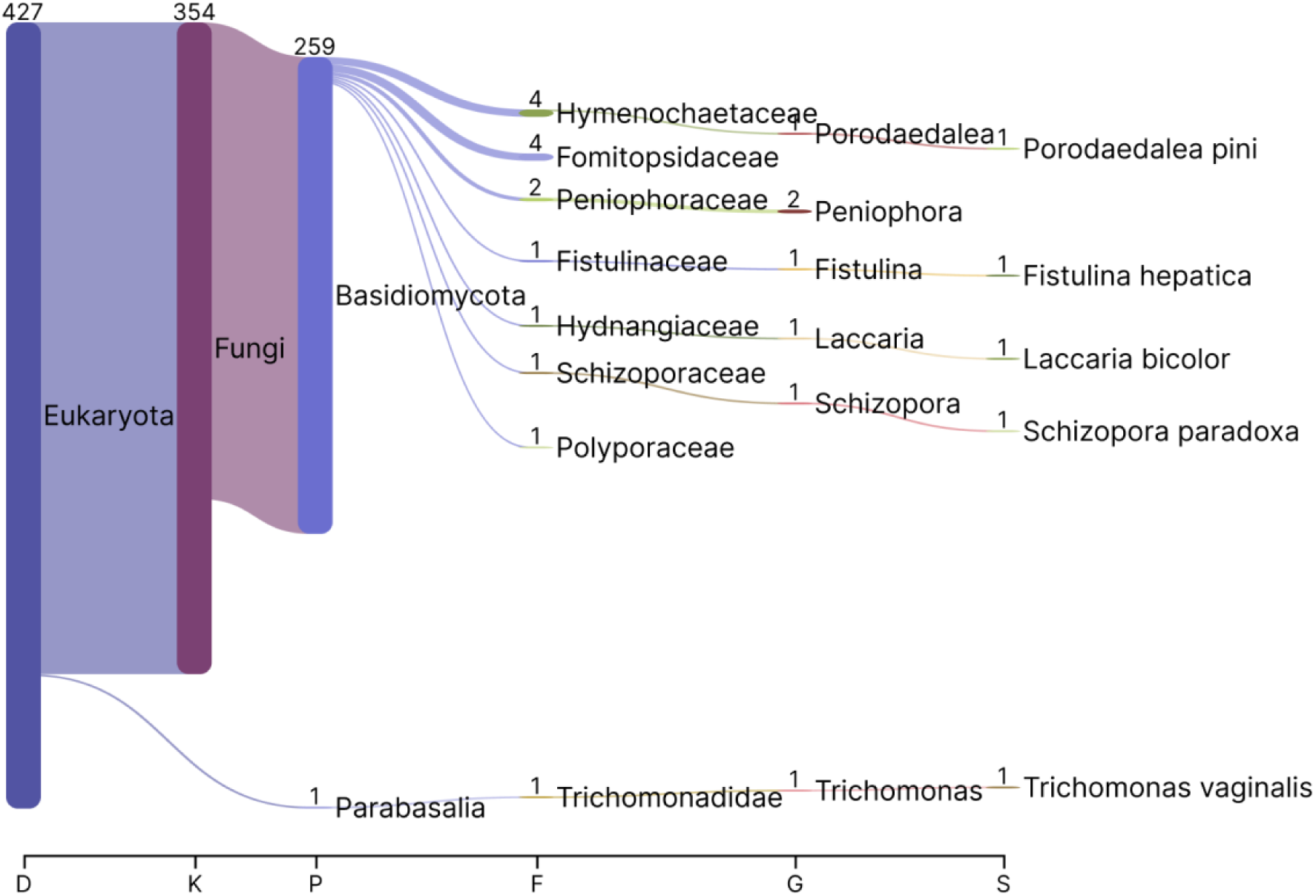
The taxonomic distribution of the clusters specific to the biome: Air.

**Supplementary Figure 3.**
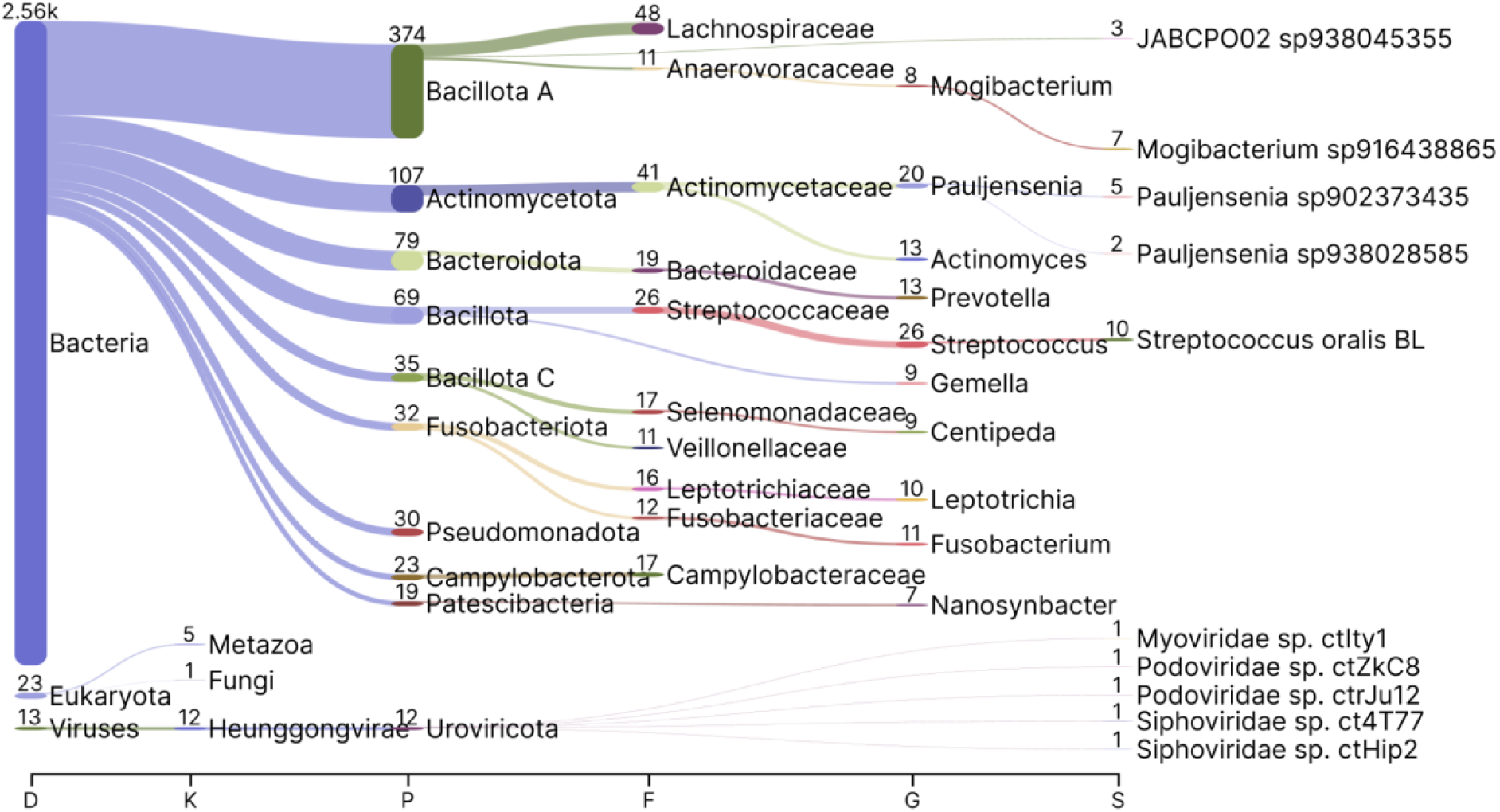
The taxonomic distribution of the clusters specific to the biome: Human digestive system.

**Supplementary Figure 4.**
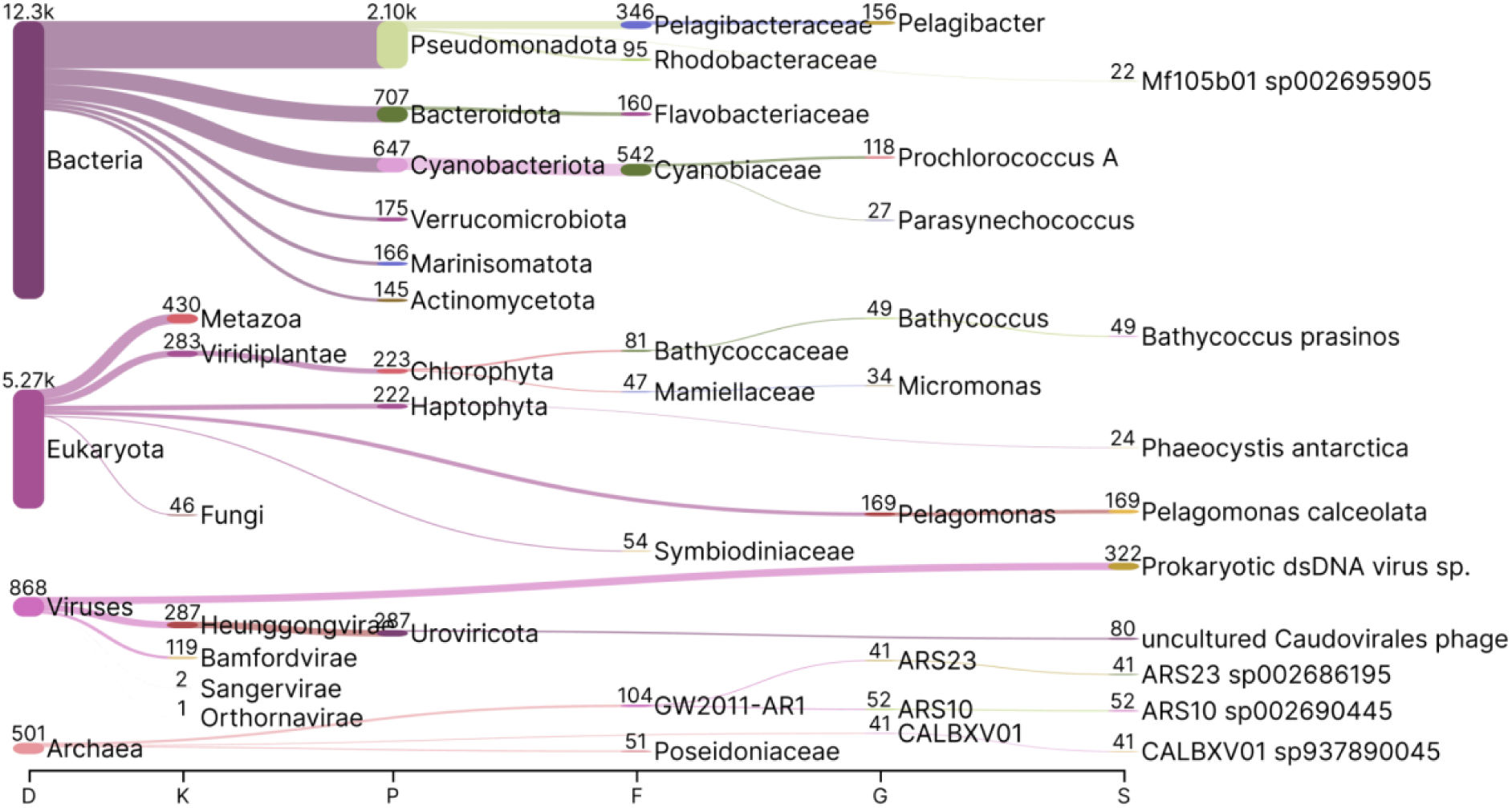
The taxonomic distribution of the clusters specific to the biome: Marine.

**Supplementary Figure 5.**
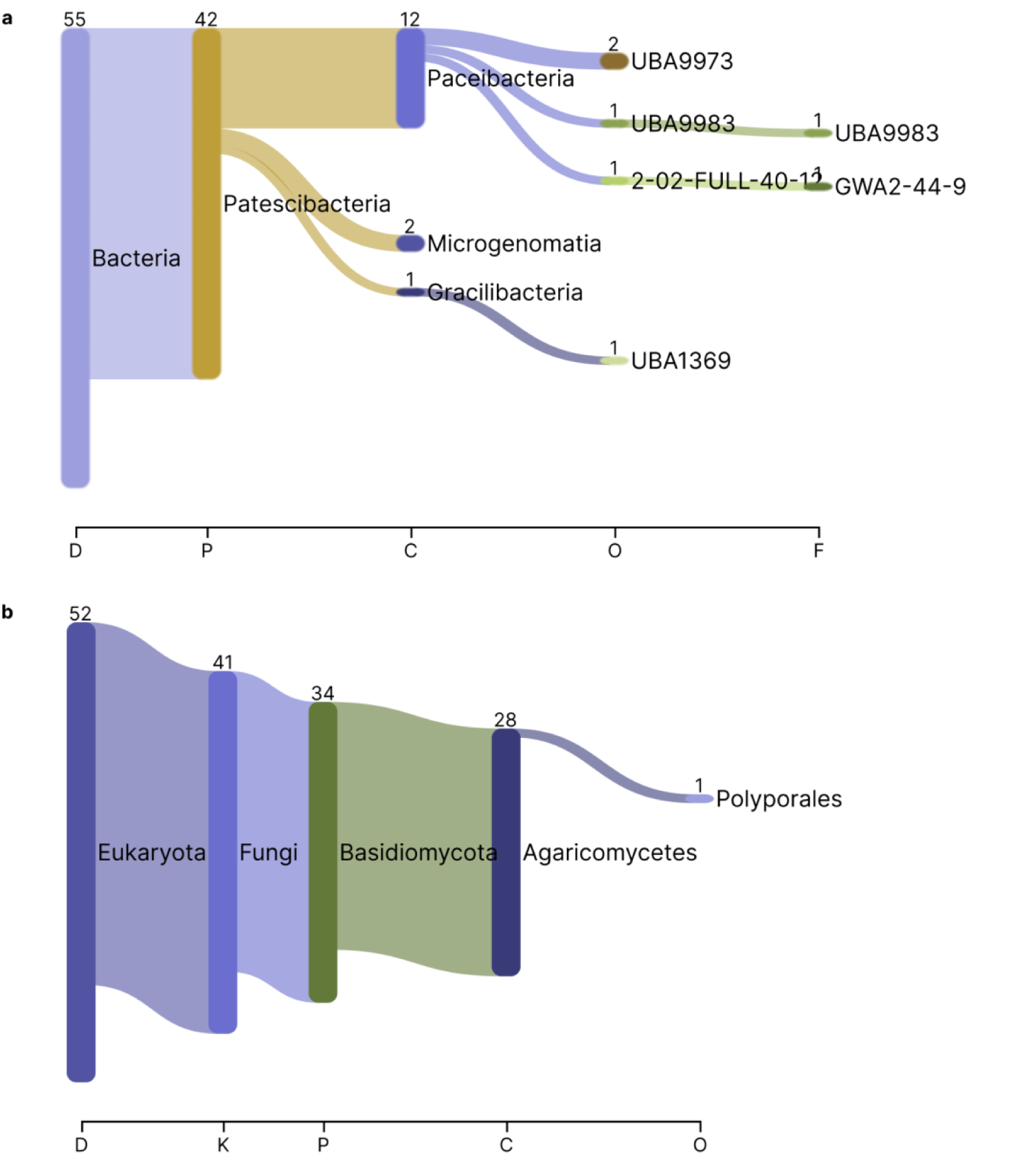
Taxonomical distribution of the LCA from the biome specific clusters with a specific Pfam. a, 69 Freshwater specific clusters that has the Type IV secretion system pilin b, 52 Air specific clusters that has the F-box domain

**Supplementary Figure 6.**
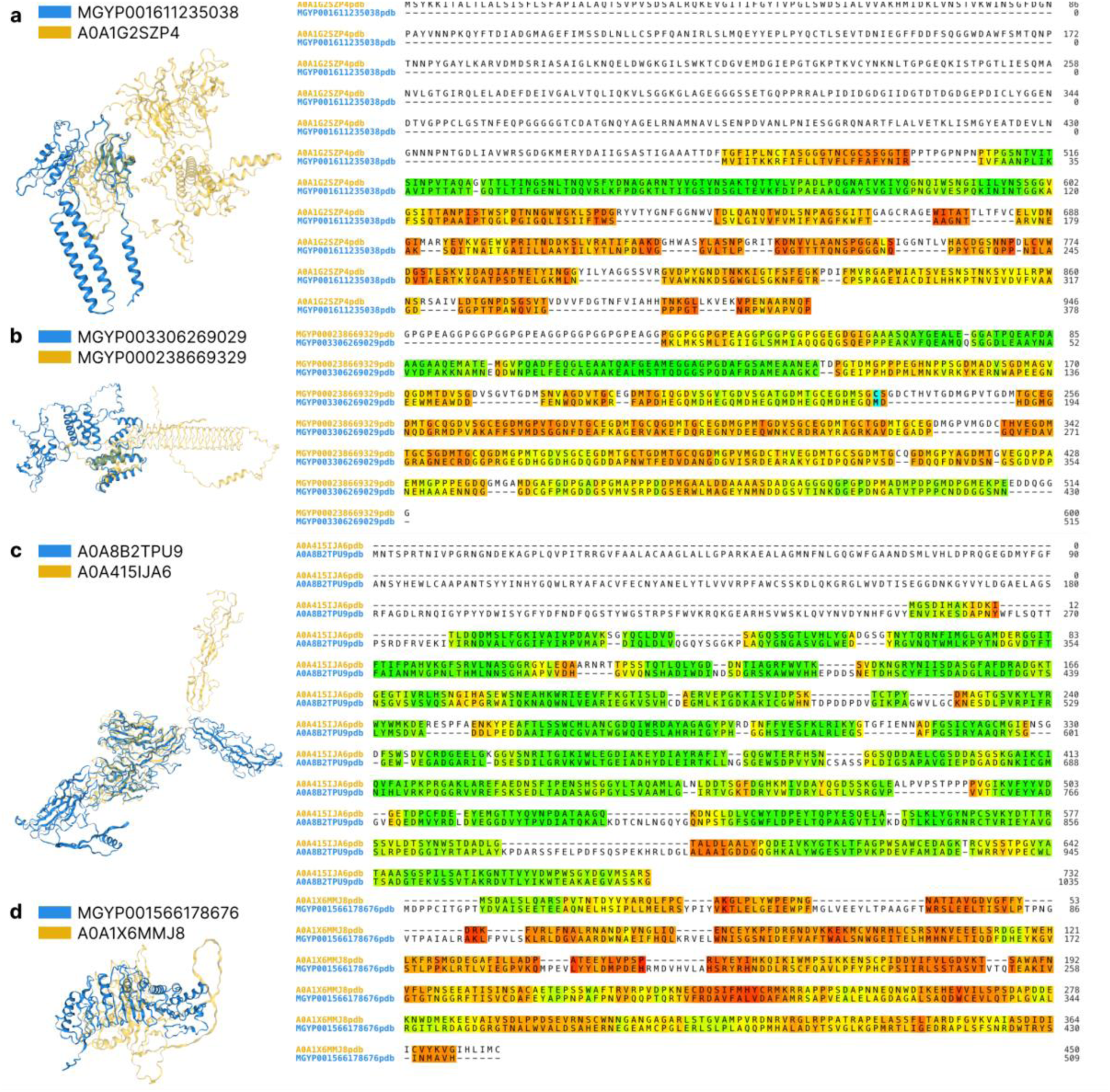
The superposition of the extreme environment specific clusters to the proteins among the non-specific clusters with the lowest E-value. The proteins - one from the environment specific clusters and the other from the non environment specific clusters - are at the left side and the lDDT scores between the proteins computed with Foldseek are at the right side. a, MGYP001611235038 (Freshwater) is aligned to A0A1G2SZP4 (Non-specific). b, MGYP003306269029 (Marine) superposed to MGYP000238669329 (Non-specific). c, A0A8B2TPU9 (Human digestive system) is superposed to A0A415IJA6 (Non-specific). d, MGYP001566178676 (Air) superposed to A0A1X6MMJ8 (Non-specific)

**Supplementary Figure 7.**
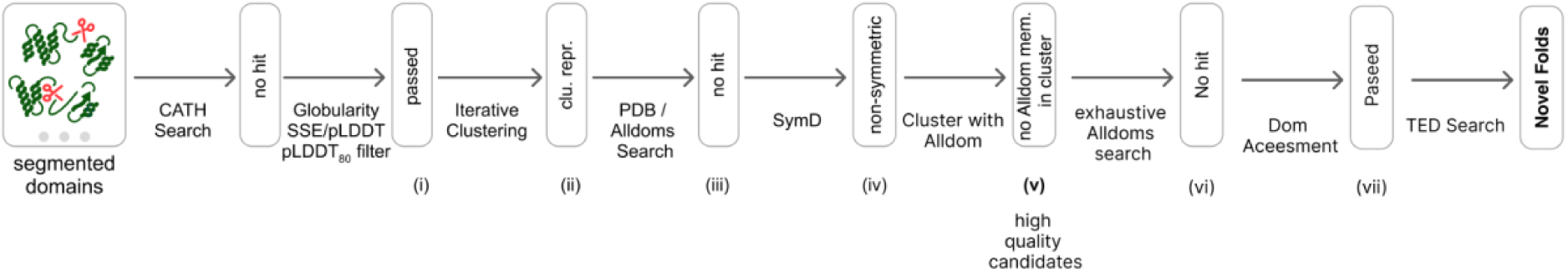
Novel fold identification workflow. Domains detected in the input protein entries that pass the quality assessment underwent iterative Foldseek clustering followed by searches against the PDB and other domain classification resources (Alldoms), with domains without matches subjected to further refinement using symmetry analysis (SymD). Remaining non-symmetric, high-confidence domains were reclustered and re-queried against Alldoms, and only those with no structural matches advanced to a final stage involving exhaustive searches against Alldoms and TED together with domain-boundary assessment. This process yielded the subset of domains that could not be assigned to any known fold in existing structural classifications.

**Supplementary Table 1.** Distribution of predicted protein domains across confidence bins Each row represents a dataset (coloured) or control bins and columns i–vii correspond to each step in the Novel fold identification defined in Supplementary Figure 7, showing the number of domains survived in each step. Step v (termed “high-quality candidates”) was chosen as the comparison point between the controls and the focus fraction, because it represents the last stage at which a substantial number of candidates remain, providing sufficient statistical power to detect differences in novelty potential between categories. The 10,000-entry control samples from the low-confidence and singleton categories were drawn uniformly at random from their respective pools of cluster representatives to test the assumption that these categories have lower novelty potential (Methods).

| Datasets and control samples / pipeline step (Supp. Fig. 7) | i | ii | iii | iv | v (high-quality candidate) | vi | vii | Novel Folds |
| --- | --- | --- | --- | --- | --- | --- | --- | --- |
| Focus fraction: 0.18M high-quality no CATH match domains from the 3.2M ESM-only clusters | 183,726 | 64,697 | 15,956 | 7,461 | 3,473 | 801 | 439 | 12 |
| Focus fraction: 2.3M low-quality no CATH match domains from the 3.2M ESM-only clusters | 98,453 | 54,834 | 18,109 | 9,373 | 4,639 | 1267 | 61 | 33 |
| Control: 9548 domains from the 10K sample of low-quality sequence level cluster representatives (pLDDT<60) | 23 | 22 | 1 | 0 | 0 | 0 | 0 | 0 |
| Control: 6324 domains from the 10K sample of structural singletons | 249 | 232 | 12 | 6 | 1 | 0 | 0 | 0 |

**Supplementary Figure 8.**
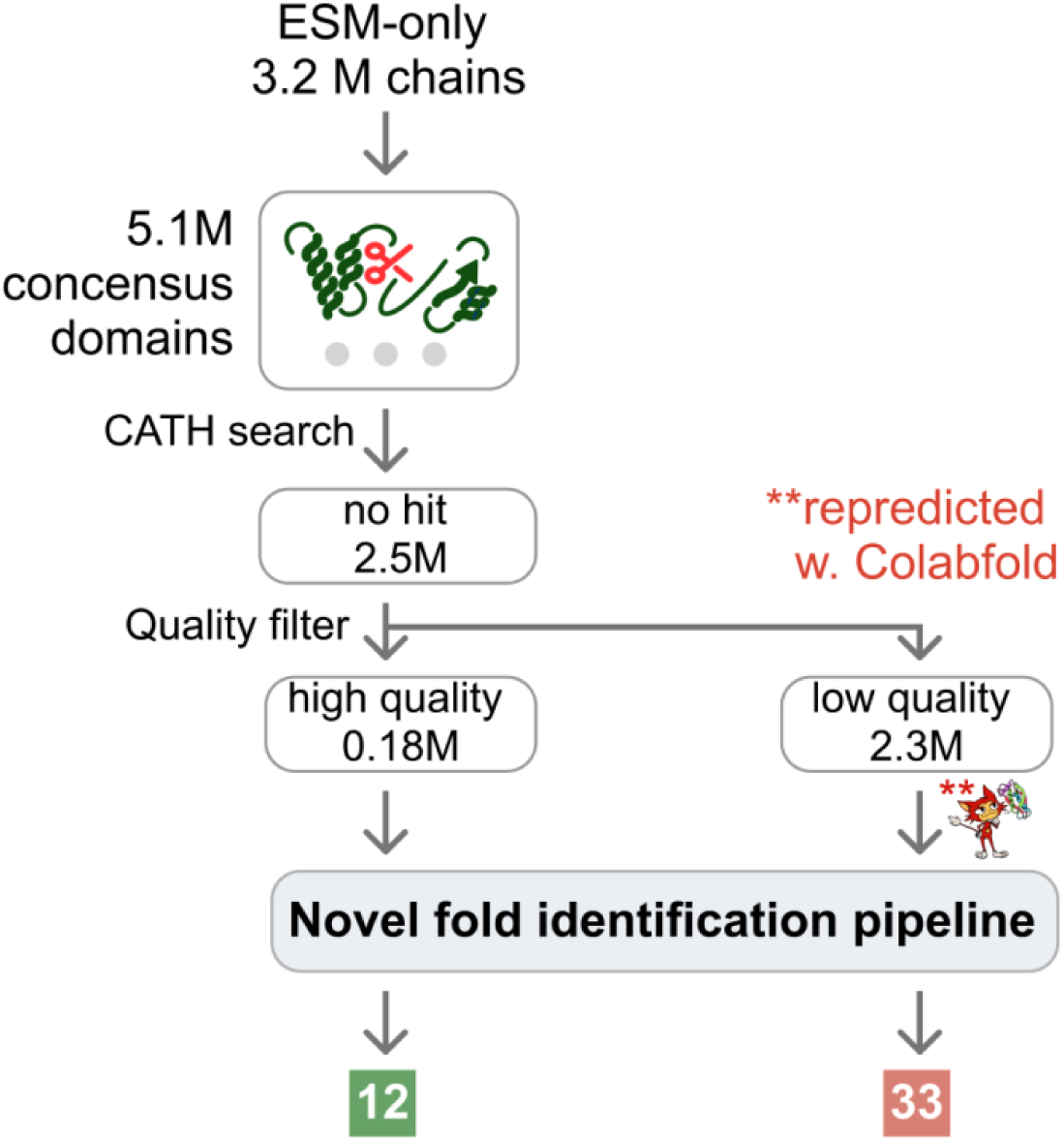
Domain-level novel fold discovery in the ESM-only non-singleton fraction. From the 5.1 million domains derived from the 3.2 million ESM-only non-singleton structural cluster representatives, 0.18 million lacked CATH assignments and passed quality filters, yielding 12 novel folds. An additional 2.3 million low-confidence domains without CATH hits were re-predicted using ColabFold, rescuing 0.42 million domains (pLDDT ≥ 70) and identifying 33 additional novel folds.

**Supplementary Figure 9.**
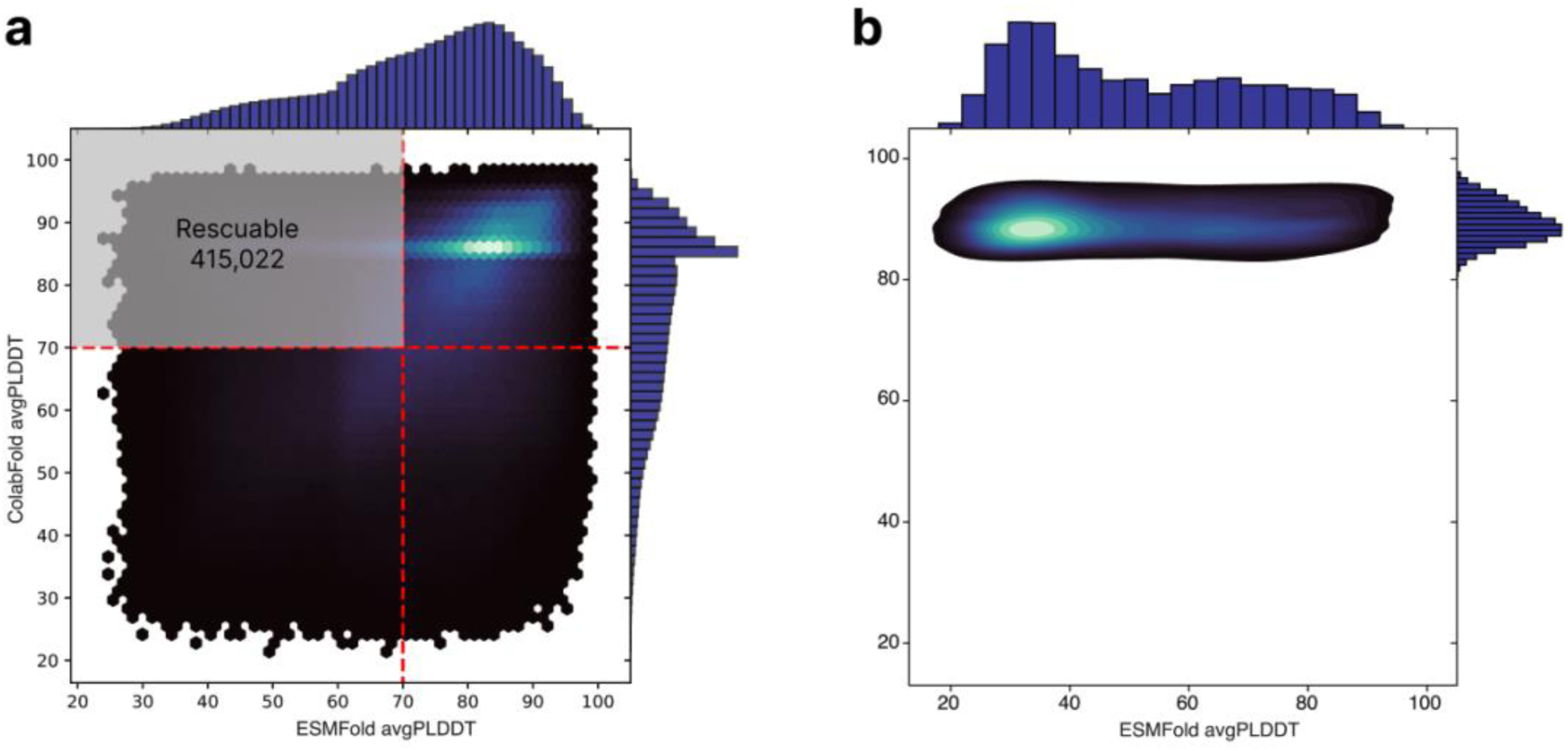
a, Density plot of average pLDDT derived from AF2 versus ESMFold for novel folds in the TED database. b, The comparison of pLDDT between the novel domains predicted by ColabFold (y-axis) and ESMFold (x-axis).

**Supplementary Figure10.**
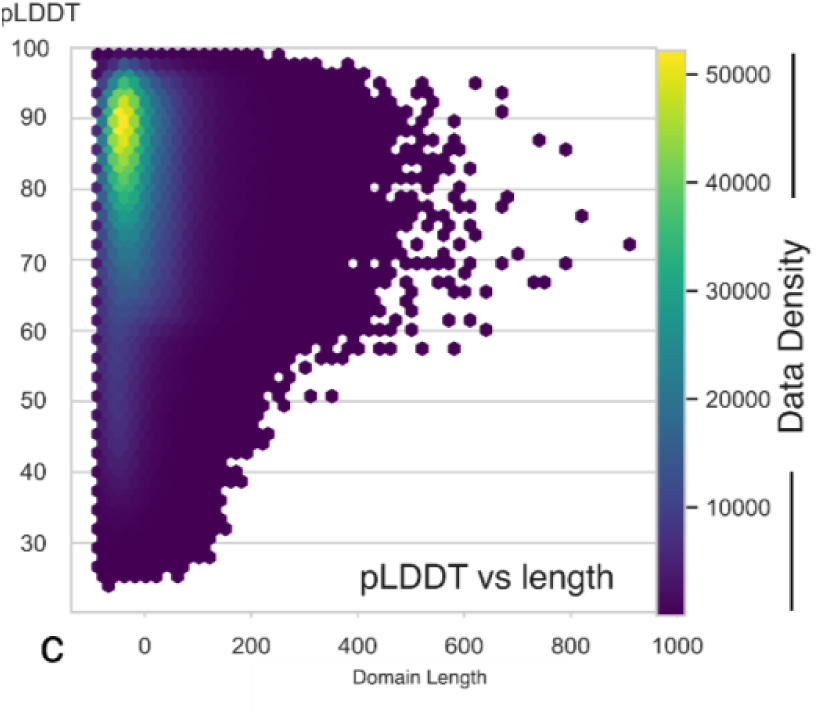
Density plot describing pLDDT as a function of domain length.

**Supplementary Figure 11.**
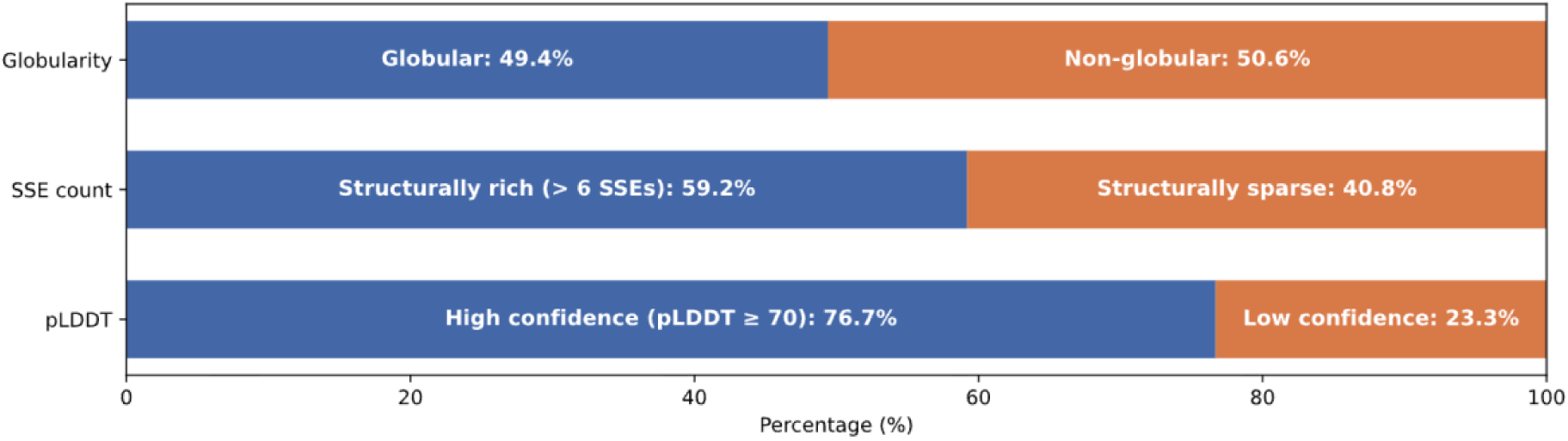
Quality assessment of domains in ESM-only clusters representatives by pLDDT, number of secondary structure elements and globularity.

Supplementary Table 2. List of 2,906 CATH H-level categories compared between Novel and Non-novel MDPs. Each row includes the CATH identifier, the presence and number of occurrences in the Novel and Non-novel sets, the type of statistical test used (Chi-square test or Fisher’s exact test) to assess over- or underrepresentation, p-value, Benjamini–Hochberg adjusted p-value, the log₂ ratio of frequencies, the differential abundance label (overrepresented or underrepresented), the number of unique novel domain pairing partners, and the name of the CATH H-level category.

The table is provided as a separate tsv file at: https://afesm.steineggerlab.workers.dev/ File name: 6-cathID_presenceFlag_nNovel_nNonNovel_statTestMethod_pValue_adjPValue_log2Ratio_d diffAbundanceFlag_nNovelPartners_cathName.tsv

**Supplementary Table 3.** List of 10 most overrepresented CATH H-level categories with significant differential abundance between Novel and Non-novel MDPs, sorted by unique number of novel domain pairing partners. Membrane-associated domains are shown in bold. . For each domain, the table includes the CATH identifier, number of occurrences in both sets, log₂(freq. in Novel set / freq. in Non-novel set), adjusted p-value (Benjamini–Hochberg corrected), and the number of unique novel domain partners.

| Category | cathname | log2ratio (x) | -log10p_adj (y) | nNovelPair |
| --- | --- | --- | --- | --- |
| 2.170.130.10 | **TonB-dependent receptor, plug domain** | 2.2807 | 60.7733 | 115 |
| 3.65.10.10 | Enolpyruvate transferase domain | 3.7650 | 160.8717 | 103 |
| 3.30.2090.10 | **Multidrug efflux transporter AcrB TolC docking domain; DN and DC subdomains** | 2.9292 | 75.2153 | 83 |
| 1.10.3720.10 | **MetI-like** | 2.8088 | 66.9116 | 82 |
| 3.40.1160.10 | Acetylglutamate kinase-like | 2.7435 | 61.8651 | 82 |
| 2.160.10.10 | Hexapeptide repeat proteins | 1.5259 | 17.1761 | 81 |
| 2.70.150.10 | **Calcium-transporting ATPase, cytoplasmic transduction domain A** | 2.6197 | 58.1307 | 81 |
| 3.40.120.10 | Alpha-D-Glucose-1,6-Bisphosphate, subunit A, domain 3 | 2.4824 | 50.2820 | 81 |
| 3.90.226.10 | 2-enoyl-CoA Hydratase; Chain A, domain 1 | 1.4943 | 17.5444 | 79 |
| 3.40.710.10 | DD-peptidase/beta-lactamase superfamily | 1.4637 | 16.5869 | 77 |

**Supplementary Figure 12.**
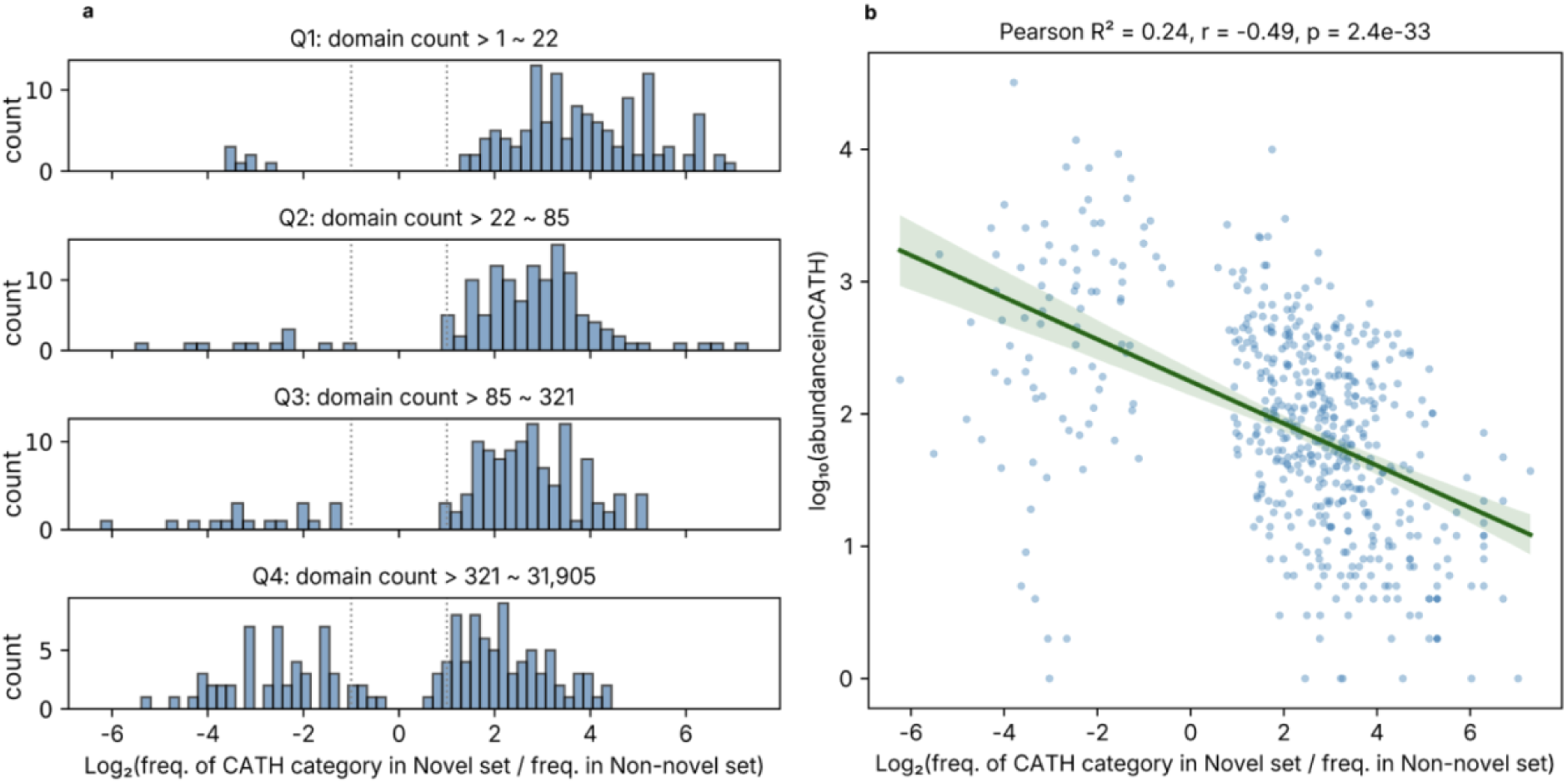
a, Distribution of log₂(Frequency of CATH H-level categories from Novel set/Frequency of CATH H-level categories from Non-Novel), across quantiles of domain counts in CATH b, Negative correlation between log₂ fold change (log₂FC) and log₁₀-transformed abundance of CATH domains. Each dot represents a single protein domain. A linear regression line is shown in green. The Pearson correlation coefficient indicates a moderate negative correlation (r = –0.49, R^2^ = 0.24), which is statistically significant (p = 2.4 × 10^−33^), suggesting that lower-abundance domains tend to show higher fold changes in expression.

**Supplementary Table 4.** Top 10 CATH H-level domains by the number of their unique novel pairing partners, ranked by log₂(freq. in Novel set / freq. in Non-novel set). Each entry includes the domain identifier, enrichment score (log₂ ratio), adjusted p-value, and the number of unique novel domain partners. These domains feature prominently in Fig. 4e as part of unexpected or functionally intriguing domain combinations.

| Category | cathname | log2ratio (x) | -log10p_adj (y) | nNovelPair | domain_count_CATHDB |
| --- | --- | --- | --- | --- | --- |
| 3.40.1570.10 | HemS/ChuS/ChuX like domains | 7.2910 | 6.0082 | 5 | 37 |
| 3.50.90.10 | YerB-like | 7.0280 | 22.2711 | 22 | 1 |
| 1.10.700.10 | Dioxygenase LigAB, LigA subunit | 6.7061 | 3.3507 | 3 | 22 |
| 3.30.500.20 | BH3703-like domains | 6.7061 | 3.3507 | 4 | 4 |
| 3.50.70.10 | - | 6.7061 | 3.3507 | 4 | 47 |
| 1.10.3480.10 | TorD-like | 6.2910 | 5.3038 | 5 | 22 |
| 1.10.4010.10 | Type II deoxyuridine triphosphatase | 6.2910 | 2.0444 | 2 | 8 |
| 1.20.1500.10 | YheA/YmcA-like | 6.2910 | 2.0444 | 2 | 12 |
| 2.30.170.40 | Ribosomal protein L28/L24 | 6.2910 | 2.0444 | 2 | 72 |
| 3.10.640.10 | Restriction endonuclease-like alpha-beta roll domain | 6.2910 | 2.0444 | 2 | 5 |

**Supplementary Figure 13.**
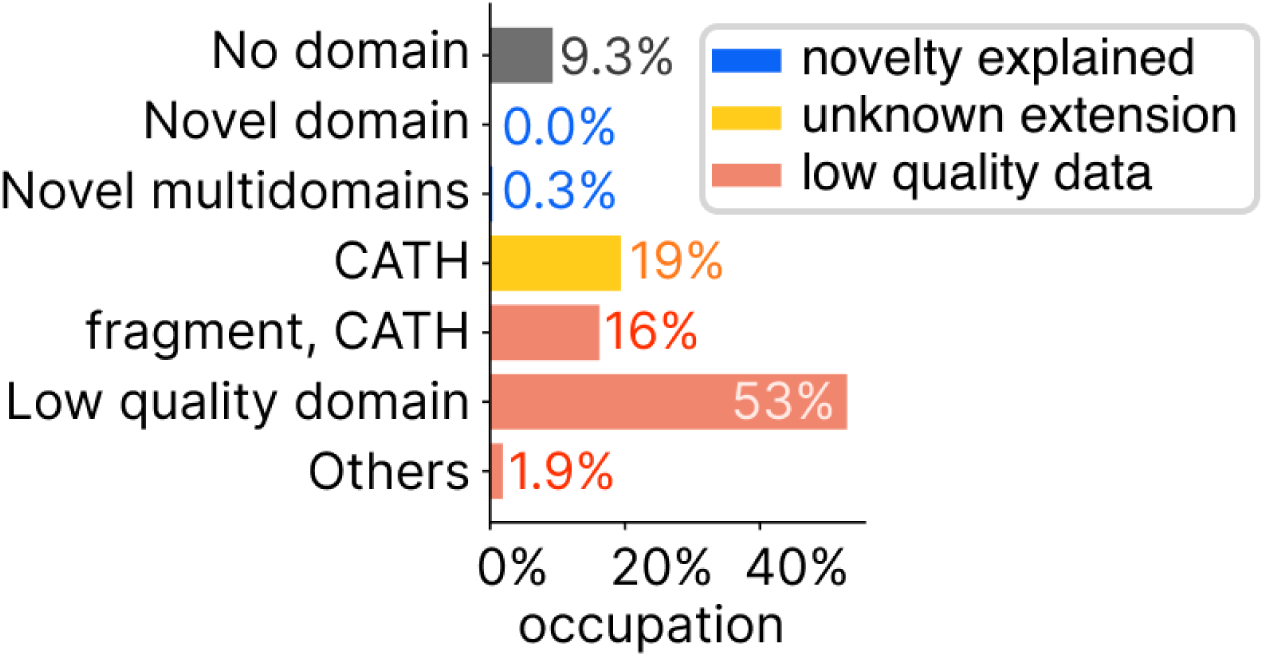
The description of the ESM-only representatives: No domain identified (9.3%), (1) Novelty explained - structurally novel domains (0.0%) or novel multi-domain arrangements (0.3%), (2) Unknown extensions - CATH domains are identified (19%) potentially containing additional regions, (3) Low-quality data - fragments (16%) or predicted domains filtered due to the quality filtering (53%) or other criteria (annotatable domains by iterative clustering and search against PDB or other domain databases, symmetry and quality assessment by Foldclass, 1.9%).

## Notes

https://afesm.foldseek.com/

### Summary of Updates

We raised the manuscript according to reviewer feedback

